# SIRT5 acts in the tumor microenvironment via endothelial cell metabolism to support breast cancer growth

**DOI:** 10.64898/2026.04.09.717584

**Authors:** Anthony M. Chen, Issahy Cano, Qian Zhao, Pei-Yin Tsai, Elijah A. Bacchus, Sadhan Jana, Irma R. Fernandez, Zeribe C. Nwosu, Andrew Miller, Joeva Barrow, Hening Lin, Esak Lee, Robert S. Weiss

## Abstract

Triple-negative breast cancer (TNBC) is characterized by aggressive progression and poor prognosis, partly due to abnormal angiogenesis. While the metabolic reprogramming of tumor cells is well characterized, the metabolic regulation of tumor-associated endothelial cells (ECs) remains unclear. Here, we identified the mitochondrial deacylase SIRT5, which has established tumor-promoting roles in TNBC cells, as a key regulator of endothelial metabolic homeostasis and tumor angiogenesis. SIRT5-deficient host mice showed significant defects in supporting the growth of orthotopic SIRT5-proficient mammary tumor transplants, and the resulting neoplasms showed defects in tumor vascularization. In a 3D microfluidic vessel-on-chip model, *SIRT5* loss compromised vascular barrier integrity and EC sprouting. Mechanistically, *SIRT5*-deficient ECs exhibited diminished mitochondrial respiratory capacity but apparently normal glycolysis. SIRT5 loss also caused increased mitochondrial reactive oxygen species levels, and a mitochondrial antioxidant rescued the endothelial cell defects following SIRT5 loss, indicating that SIRT5-mediated mitochondrial redox homeostasis in the tumor microenvironment is necessary to maintain vascular function. Orthotopic co-transplantation of TNBC and EC cells with or without SIRT5 knockdown demonstrated that endothelial SIRT5 promotes increased tumor growth *in vivo*. These results suggest that targeting SIRT5 offers a potential therapeutic strategy to disrupt tumor angiogenesis and suppress TNBC progression by targeting the metabolic vulnerabilities of the tumor endothelium.

## INTRODUCTION

Breast cancer (BC) is a leading cause of cancer-related death in women worldwide ^1^. Triple-negative breast cancer (TNBC) accounts for 15–20% of cases and is characterized by aggressive progression and poor prognosis due to a lack of targeted therapies ^2^. While treatments such as chemotherapy exist, recurrence and drug resistance often develop due to the heterogeneity of TNBC, making treatments that target the tumor microenvironment (TME) a promising alternative strategy. The abnormal tumor vasculature that supports rapid TNBC growth represents an important therapeutic target ^3^. Tumor angiogenesis allows rapid growth and metastasis but results in structurally abnormal vessels that exacerbate hypoxia and limit drug delivery ^4^. Although there are FDA-approved anti-angiogenic therapies targeting vascular endothelial growth factor (VEGF) ^5^, an angiogenic factor, they yield only modest survival benefits as a monotherapy due to resistance and compensatory signaling ^6–8^. This highlights the need to target endothelial cells (ECs) that receive angiogenic signaling, as their genetic stability makes them less susceptible to drug resistance than highly heterogeneous cancer cells, which can develop resistance through secreting various angiogenic factors and other mechanisms.

In the TME, endothelial metabolism is reprogrammed to support rapid sprouting. Tumor ECs upregulate glycolysis—a “Warburg-like” metabolism to generate ATP under hypoxia while diverting glucose carbons into biosynthetic pathways ^9–12^. While tumor ECs rely on glycolysis for energy, endothelial mitochondria function as essential biosynthetic and signaling organelles ^13^. Mitochondrial reactive oxygen species (mtROS) act as signaling messengers that promote angiogenic sprouting ^14^. However, excessive ROS causes oxidative damage and barrier dysfunction ^15,16^. Therefore, tumor ECs must tightly regulate mitochondrial redox balance to survive the stressful TME.

The mitochondrial sirtuin SIRT5 is a key regulator of mitochondrial metabolism and redox homeostasis. Uniquely among the mitochondrial sirtuins, SIRT5 functions as a lysine desuccinylase, demalonylase, and deglutarylase, modulating enzymes regulating fatty acid oxidation, the TCA cycle, and ROS detoxification ^17–19^. SIRT5 has context-dependent roles in cancer, functioning as a tumor suppressor in some cancers but as a tumor promoter in others ^20–24^. In breast cancer, SIRT5 is frequently amplified and overexpressed, particularly in basal breast cancers, and SIRT5 inactivation suppresses the transformed properties of cultured breast cancer and impairs tumor progression and metastasis in mouse models ^22,23^. Mice lacking *Sirt5* or treated with a SIRT5-selective inhibitor show minimal side effects ^22,25^, highlighting the potential of SIRT5 as a safe target for cancer therapies.

Despite important roles for SIRT5 in cancer cells, its function in the TME remains unknown. Here, we identify SIRT5 as an important metabolic regulator in ECs that protects mitochondrial homeostasis under the stressful conditions of tumor angiogenesis. We show that SIRT5 loss results in impaired mitochondrial respiration and pathological elevation of mtROS. This redox imbalance compromises endothelial barrier function, leading to the formation of structurally defective, leaky vessels that are unable to sustain perfusion. Consequently, SIRT5-deficient EC cells fail to support primary tumor outgrowth and lung metastatic dissemination in a xenograft TNBC model, establishing SIRT5 inhibition as a strategy to inhibit the vascular support system of aggressive BCs.

## METHODS

### Mouse breeding and husbandry

All animal experiments were performed under an approved IACUC protocol and in compliance with institutional and federal guidelines. Animals were maintained in an AAALAC-accredited facility with veterinary oversight. *Sirt5* knockout (KO) mice were backcrossed with wild-type (WT) C57BL/6 mice for more than 10 generations ^24^. *Sirt5*^+/−^ mice were interbred to produce *Sirt5*^+/+^ (WT) and *Sirt5*^-/-^ (KO) mice. B6.FVB-Tg(MMTV-PyVT)634Mul/LellJ (PyMT) transgenic mice were obtained from The Jackson Laboratory ^26^. Experimental mice carrying the PyMT transgene were generated by crossing *Sirt5*^+/−^ female mice to *Sirt5*^+/−^ PyMT-positive male breeders. All PyMT mice were collected 50 days after tumor detection by palpation.

### Cell culture

Human umbilical vein endothelial cells (HUVECs) and MDA-MB-231 cells were purchased from ATCC. AT-3 mouse mammary carcinoma cells were acquired from S. Abrams (Roswell Park Comprehensive Cancer Center) ^27^. HUVECs were maintained in Human Endothelial Cell Growth Medium V2 (ECGM, PromoCell, 213-500). Cells were incubated at 37°C in a humidified atmosphere containing 5% CO₂ and passaged at 80–90% confluence using 0.05% trypsin-EDTA (Gibco, 15400054). Only early-passage cells (P3–P8) were used for experiments. SIRT5 knockdown (KD) HUVEC cells were generated by transduction with lentiviral particles encoding either shSIRT5-1 (TRCN0000232660), shSIRT5-2 (TRCN0000232661), or a non-targeting control shSC (SHC202). Lentiviral production was performed using Lipofectamine 3000 (Invitrogen, L3000015) following the guide provided by Invitrogen, and successfully transduced cells were selected and expanded for downstream assays. MDA-MB-231 cells were maintained in RPMI 1640 medium (Gibco, 11875093) with 10% bovine calf serum (BCS, HyClone, SH30072.03). AT-3 cells ^28^ were maintained in Dulbecco’s Modified Eagle Medium (DMEM, Gibco, 11005065) with 10% BCS.

### Tumor Transplant Mouse Models

For syngeneic transplant experiments, a total of 2 × 10⁵ AT-3 murine mammary carcinoma cells were resuspended in 20 µL of a 1:1 mixture of PBS and growth factor-reduced Matrigel (Corning, 354234) and injected into the inguinal mammary fat pads of 5–6-week-old *Sirt5* WT and KO female mice. For co-implantation xenograft experiments, 3 × 10⁵ MDA-MB-231 cells and 2 × 10⁶ HUVECs were resuspended in 10 µL of PBS, respectively. The cell suspensions were mixed and injected into the inguinal (#4) mammary fat pads of 5–6-week-old NOD.Cg-*Prkdc*ˢᶜⁱᵈ *Il2rg*ᵗᵐ¹ᵂʲˡ/SzJ (NSG) female mice. Tumors are collected based on the humane endpoint or time endpoint specified in each experiment.

### Tumor Tissue Processing

For paraffin embedding, excised tumors were fixed in 4% paraformaldehyde (PFA) at room temperature for 24 hours. For optimal cutting temperature (OCT) compound embedding, tumors were fixed in 4% PFA at 4°C for 4 hours, incubated in 30% sucrose (w/v) in PBS at 4°C overnight, and then embedded in OCT compound and stored at −80°C until sectioning.

### Immunohistochemistry (IHC) Staining

5 µm thick formalin-fixed paraffin-embedded (FFPE) tissue sections on glass slides were deparaffinized and rehydrated with graded alcohols. Antigen retrieval was performed by microwaving the slides for 20 minutes in citrate buffer (pH 6.0) or Tris-EDTA buffer (10 mM Tris, 1 mM EDTA, pH 9.0). Slides were incubated with 3% hydrogen peroxide in methanol for 20 minutes at room temperature to inhibit endogenous peroxidase activity. Slides were blocked with TBS-T containing 4% BSA and 5% goat serum and incubated with primary antibodies overnight at 4°C. Slides were then incubated with biotinylated secondary antibody, followed by streptavidin-HRP conjugate (Vector Labs, SA-5704-100) at room temperature for 10 minutes. Immunoreactivity was visualized with DAB (Thermo Fisher, 34002), counterstained with hematoxylin (Ricca, 3536-32), dehydrated, and mounted. Slides were scanned using an Aperio ScanScope and analyzed using ImageJ.

### Western blot

Cell lysates were prepared using lysis buffer (10% glycerol, 1% Triton X-100, 1mM PMSF, 21 µM Leupeptin, 1 mM sodium orthovanadate, 1 mM EGTA, 10 mM NaF, 1 mM Sodium pyrophosphate, 1 mM ß-glycerophosphate, 20 mM Tris-HCL, 137 mM sodium chloride, 5 mM EDTA, and 5 mM nicotinamide in double-distilled water). Lysates were then centrifuged at 17,000 x g for 20 minutes at 4°C to remove the cell debris. Protein concentration was measured with the Bradford method. 20–40 ug of protein were separated with a 10% polyacrylamide gel, and transferred to a PVDF membrane (IPVH00010, Millipore). The membrane was blocked in TBST (25 mM Tris-HCl, pH 7.4, 150 mM NaCl, and 0.1% Tween-20) containing 3% BSA and, incubated with antibodies in TBST containing 3% BSA, and incubated with secondary antibodies in TBST. The chemiluminescent signal was detected with ChemiDoc XRS Molecular Imager (Bio-Rad).

### Proliferation assay

HUVECs were seeded into 96-well plates at 5,000 cells per well in 200 μL ECGM and incubated at 37°C in a humidified 5% CO₂ atmosphere. At each time point (24, 48, and 72 hours post-seeding), the CyQUANT Direct Proliferation Assay Kit (Thermo Fisher, C25011) was used to measure proliferation according to the manufacturer’s protocol.

### Trans-well migration/invasion assay

HUVECs were seeded into trans-wells (Corning® FluoroBlok™, Corning, 351152) in 24-well plates with Human EC Basal Medium (ECBM, PromoCell, 210-500) supplemented with 0.5% fetal bovine serum (FBS, A5256701, Gibco) (25,000 cells/300 ul/well). For the invasion assay, Matrigel (Corning, 354234) was diluted to 0.3 mg/ml with ECBM, and 100 μl were added to the trans-wells and incubated at 37°C in a humidified 5% CO₂ atmosphere for 1 hour before cells were seeded. 700 μl ECGM was added to the bottom chamber after cells were seeded to induce migration/invasion. After incubation at 37°C in a humidified 5% CO₂ atmosphere for 24 hours, cells were fixed with 4% PFA for 10 minutes and stained with 1 μg/ml propidium iodide (Thermo Fisher, P3566) for 15 minutes for fluorescence imaging to quantify the number of cells that migrated/invaded.

### Tube formation assay

Matrigel (Corning, 354234) was added to 96-well plates (100 μl/well) and incubated at 37°C in a humidified 5% CO₂ atmosphere for 1 hour before cell seeding. HUVECs were seeded into the Matrigel-coated 96-well plate with ECGM (10,000 cells/100 ul/well) and incubated at 37°C in a humidified 5% CO₂ atmosphere for 24 hours. The tubular structure formed by HUVECs was imaged. The lengths of the tubular structure’s segments were measured with ImageJ.

### Drug treatment

For experiments involving the SIRT5-selective inhibitor DK1-04e^22^ (synthesized in-house) or MitoTEMPO (Sigma-Aldrich, SML0737), cells were pre-treated with the indicated compound for 24 hours prior to the start of the assay. Dimethyl sulfoxide (DMSO, Sigma-Aldrich, D2650) was used as the vehicle for both drug treatments. Drug treatments were maintained throughout the duration of the experiment and refreshed every 24 hours unless otherwise specified.

### Vessel-on-chip

Microfluidic chip fabrication was performed as previously described ^29,30^. Briefly, polydimethylsiloxane (PDMS) was poured into a mold, cured, and cut to shape. The PDMS chips and glass coverslips were then plasma-treated and assembled to form sealed devices. To facilitate cell adhesion, the chips were coated with poly-L-lysine, followed by the addition of collagen in the presence of needles to create defined chamber spaces. After collagen polymerization, the needles were carefully removed, and HUVECs were seeded at a density of 1,000,000 cells per chip for subsequent experiments. Culture media were refreshed daily. After the desired live-cell procedures, cells were fixed with 4% paraformaldehyde (PFA), immunostained, and imaged using Leica SP8 Confocal Microscope and Leica Application Suite X Office version 3.5.5.19976. Z-stack images were acquired and processed using maximum projection for further analysis in ImageJ.

### Permeability assay

Culture media were removed from the microfluidic chips, and fluorescence-labeled 70 kDa dextran (D1830, Invitrogen) in ECBM was added to the chamber containing HUVECs. Time-lapse images (time point X = t_x_) were acquired using a confocal microscope and analyzed with a MATLAB-based algorithm to quantify diffusive permeability and vessel diameter, as previously described^30^.

### Seahorse

Oxygen consumption rate (OCR) and extracellular acidification rate (ECAR) were measured with Agilent Seahorse XFe Flux Analyzer (Agilent Technologies) to assess metabolic functions as described previously ^31,32^. HUVECs were seeded in 24-well Seahorse XFe24 cell culture microplates at a density of 20,000 cells per well with ECGM. On the day of the assay, cells were washed and switched to unbuffered DMEM supplemented with 4.5 g/L glucose, 4 mM glutamine, and 100 mM sodium pyruvate, pH 7.4. The Mito Stress Test was performed following the user guide provided by Agilent, with drug concentrations as follows: oligomycin A (1.5 µM), FCCP (2 µM), Rotenone (1.25 µM), and Antimycin A (2.5 µM). For extracellular acidification rate (ECAR) measurements, cells were washed and switched to unbuffered DMEM supplemented with 2mM glutamine (pH 7.4) for 1 hour, following the user guide provided by Agilent. Drug concentrations for the glycolysis stress test were as follows: glucose (10 µM), oligomycin A (1.5 µM), 2-DG (50 µM). Respirometry data were collected using the Agilent Wave software v2.6.1 and exported to GraphPad Prism v10.

### Mitochondrial Superoxide Detection by Flow Cytometry

HUVECs were cultured to 80% confluence and pre-treated with rotenone at the desired concentration at 37°C in a humidified atmosphere containing 5% CO₂ for 2 hours. Following incubation, cells were harvested using 0.05% trypsin, centrifuged at 1000 rpm for 5 minutes, and resuspended in phosphate-buffered saline (PBS) containing 2% fetal bovine serum (FBS) at a concentration of 1 million cells per mL. Aliquots of 500 μL were transferred into individual Eppendorf tubes. MitoSOX red was added to each tube at a final concentration of 1 μM to assess mitochondrial superoxide levels. Samples were incubated at 37°C on a shaker for 30 minutes, then centrifuged to pellet the cells. The supernatant was discarded, and the cell pellets were washed twice with PBS containing 2% FBS. After each wash, cells were centrifuged, and the supernatant was removed. The final cell pellets were resuspended in 0.5 mL of PBS supplemented with 2% FBS and transferred to flow cytometry polystyrene test tubes (Corning, 352235). DAPI (Invitrogen, D1306) was added at a final concentration of 0.1 μg/mL immediately prior to measurement. Fluorescence was analyzed using a FACSmelody flow cytometer.

### Mitochondrial Superoxide Detection by Live-Cell Imaging

HUVECs were incubated with MitoSOX red (Invitrogen, M36008) for 30 minutes at a final concentration of 0.5 μM. Cells were then washed 3 times with PBS for 5 minutes each. Live-cell imaging was performed using a Leica SP8 Confocal Microscope and Leica Application Suite X Office version 3.5.5.19976.

### *In vivo* Assessment of Tumor Oxygenation and Vascular Leakiness

Tumor hypoxia was assessed by intraperitoneal injection of pimonidazole hydrochloride (Hypoxyprobe, HP-100mg, Hypoxyprobe, Inc) (60 mg/kg) 1 hour prior to euthanasia. Vascular leakiness was assessed by tail vein injection of fluorescence-labeled 70 kDa dextran (D1830, Invitrogen; 1 mg/mouse) 30 minutes prior to euthanasia. Hypoxic regions and dextran accumulation were subsequently visualized in sectioned tumor tissue by immunofluorescence staining or direct fluorescence microscopy, respectively.

### Immunofluorescence (IF) Staining

Slides carrying 7 µm-thick OCT-embedded frozen tissue sections were washed with TBS-T and blocked with TBS-T containing 10% goat serum. Slides were incubated with primary antibodies (including anti-Hypoxyprobe adduct antibodies for hypoxia assessment) overnight at 4°C. Slides were then incubated with fluorescent secondary antibodies and 1 µg/ml DAPI (Invitrogen, D1306) at room temperature for 1 hour. Fluorescence images were acquired using a Leica SP8 Confocal Microscope and processed using Leica Application Suite X Office version 3.5.5.19976. For leakiness assays, dextran fluorescence was visualized directly without additional antibody staining.

### Antibodies

SIRT5 rabbit monoclonal antibody (8782), HSP90 rabbit monoclonal antibody (4877), CD31 rabbit monoclonal antibody (77699, for IHC staining), and Rabbit IgG HRP-linked antibody (7074) were from Cell Signaling. Mouse CD31 rat monoclonal antibody (7388) and human CD31 rabbit monoclonal antibody (76533) were from Abcam. LYVE1 rabbit polyclonal antibody (11-034) was from AngioBio. VE-cadherin mouse monoclonal antibody (9989) was from Santa Cruz Biotechnology. Hypoxyprobe rabbit antibody (Pab2627) was from Hypoxyprobe, Inc. Alexa Fluor Secondary Antibodies were from ThermoFisher.

### Statistical Analyses

All statistical analyses were performed using GraphPad Prism version 10.0 (GraphPad Software, San Diego, CA, USA). Data are presented as mean ± standard deviation (SD) unless otherwise stated. One-way analysis of variance (ANOVA), unpaired two-tailed Student’s t-test, and two-way ANOVA were used to examine data in experiments. *p*-values of < 0.05 were reflected as significant. (\**p <* 0.05; \*\**p <* 0.01; \*\*\**p <* 0.001; \*\*\*\**p <* 0.0001).

## RESULTS

### Loss of tumor-extrinsic SIRT5 reduces tumor angiogenesis in mouse mammary carcinomas

Previous studies have characterized the role of tumor cell-intrinsic SIRT5 in BC cells ^22,23^, yet the function of tumor-extrinsic SIRT5 and its impact on mammary tumor progression remain unexplored. To address this gap, we assessed the effects of tumor-extrinsic SIRT5 via orthotopic injection of *Sirt5*-expressing mouse mammary carcinoma AT-3 cells into the inguinal mammary fat pads of *Sirt5* WT and KO host mice. In *Sirt5*-deficient hosts, tumor growth was significantly impaired, reducing tumor volume by ∼71% and weight by ∼58% at the endpoint compared with tumors from *Sirt5* WT hosts (*p <* 0.001; **Fig. 1A-C**). This indicates that SIRT5 acts in non-tumor cells to facilitate mammary tumor progression. Histopathology revealed a solid growth pattern, moderate necrosis and atypia, and high mitotic activity for tumors from both groups (**Fig. S1A**). Because prior studies involving SIRT5 inactivation in both cancer cells and the TME hinted at poor tumor vascularization ^22^, we examined the specific role of Sirt5 in tumor angiogenesis. The effect of global SIRT5 loss on tumor angiogenesis was investigated in tumors from *Sirt5* WT and KO PyMT mice, where *Sirt5* deletion slows mammary tumor growth, prolongs survival, reduces early tumor burden, and delays lung metastasis ^22^. In PyMT tumors, *Sirt5* loss reduced CD31-positive endothelial area by ∼38% (*p <* 0.05; **Fig. 1C-D**) and decreased the number and area of lumenized blood vessels by ∼74% and ∼90%, respectively (*p <* 0.001; **Fig. 1E-F**). Given that CD31 is expressed in both blood and lymphatic ECs, LYVE1, a lymphatic endothelial marker, was also assessed. LYVE1 positive area was unchanged between *Sirt5* WT and KO tumors (**Figure S1B-C**), indicating a specific defect in blood vessel formation. These findings indicate that SIRT5 acts to promote tumor vascularization.

**Figure 1.**
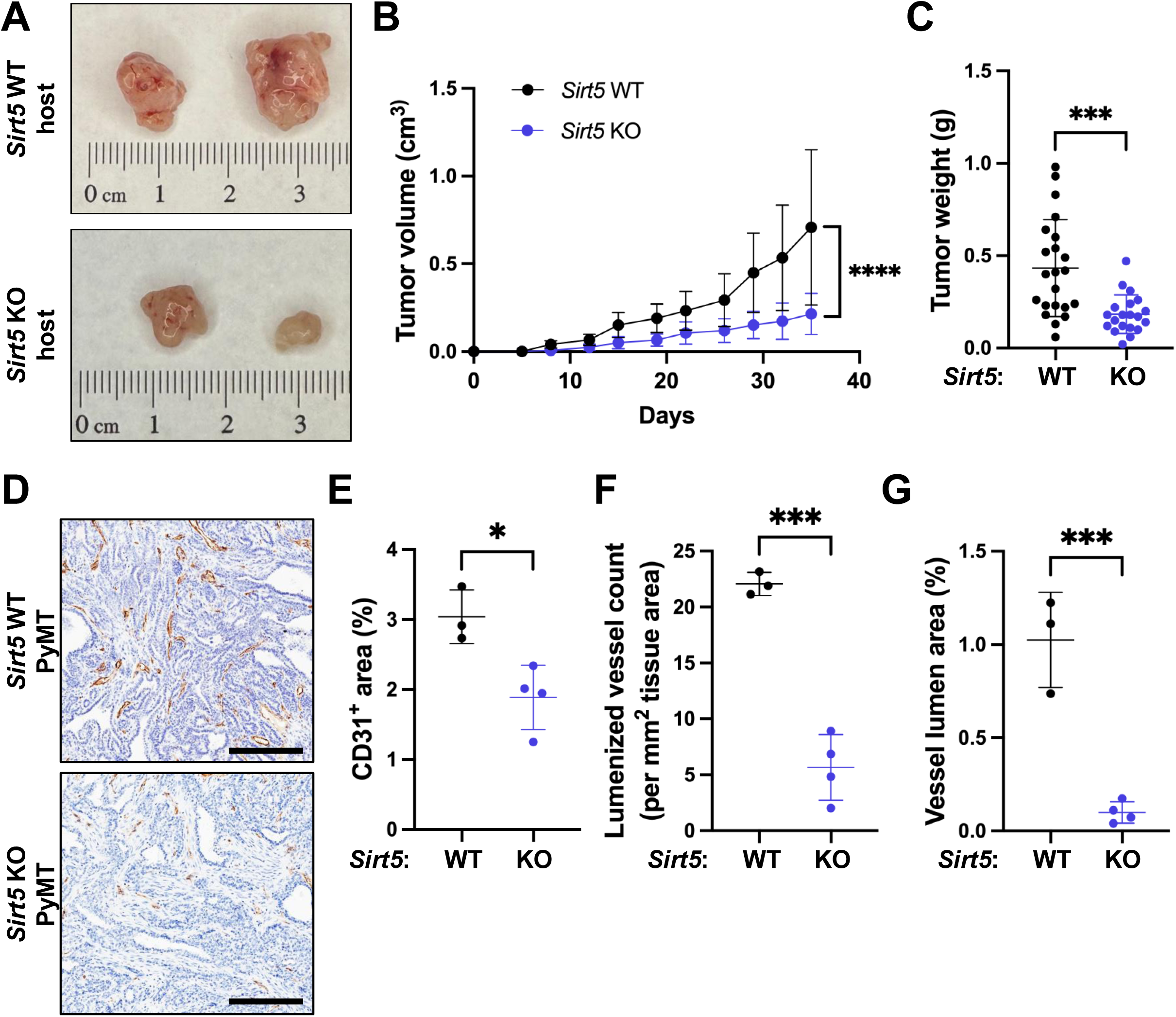
Loss of global or tumor-extrinsic *Sirt5* reduces tumor angiogenesis in mouse mammary carcinomas. A syngeneic allograft mouse model **(A-C)** and a genetically engineered mouse model **(D-G)** are used to assess the roles of SIRT5 in breast cancer progression. **(A)** Representative images of orthotopic AT3 mammary tumors in *Sirt5* WT and KO mice collected 35 days after injection. **(B-C)** Tumor growth curves and weight at endpoint of tumors from panel (A) (\*\*\**p* < 0.001, \*\*\*\**p* < 0.0001, two-way ANOVA, n = 20-22 per group). **(D)** Representative CD31 IHC images of mammary tumors from *Sirt5* WT and KO PyMT mice. Scale bar: 200 μm. **(E)** Quantification of CD31-positive area in *Sirt5* WT and KO PyMT tumors (\**P* < 0.05, unpaired two-tailed Student’s t-test, n = 3-4 per group). **(F)** Quantification of lumenized blood vessel number in *Sirt5* WT and KO PyMT tumors (\*\*\**p* < 0.001, unpaired two-tailed Student’s t-test, n = 3-4 per group). **(G)** Quantification of blood vessel lumen area in *Sirt5* WT and KO PyMT tumors (\*\*\**p* < 0.001, unpaired two-tailed Student’s t-test, n = 3-4 per group).

### SIRT5 supports endothelial cell angiogenic function in vitro

To dissect the role of endothelial SIRT5, we assessed key angiogenic functions—proliferation, migration/invasion, and tube formation—in HUVECs following *SIRT5* KD. HUVECs were transduced with lentiviral particles encoding two distinct shRNA constructs targeting *SIRT5* (shSIRT5-1 and -2) or a non-targeting scrambled control (shSC; **Fig. 2A**). Both shSIRT5 HUVECs showed decreased SIRT5 expression and elevated lysine succinylation (**Fig. 2A & Fig. S2**). *SIRT5* loss significantly impaired angiogenic functions of HUVECs. In CyQUANT direct proliferation assay, shSIRT5-1 HUVECs exhibited ∼16% and ∼20% reductions in normalized fluorescence intensity at 48 and 72 hours, respectively, while shSIRT5-2 HUVECs showed reductions of ∼33% at both time points compared with shSC HUVECs (*p <* 0.0001; **Fig. 2B**). In trans-well assays, shSIRT5-1 and shSIRT5-2 HUVECs displayed reduced migration by ∼88% and ∼73%, respectively, compared with shSC HUVECs after 24 hours (*p <* 0.001; **Fig. 2C-D**). Both *SIRT5* KD HUVECs resulted in ∼77–78% reductions in cell invasion through matrigel (*p <* 0.01; **Fig 2C&E)**. Interestingly, the reduced proliferation and invasion of *SIRT5*-deficient HUVECs echoed the *SIRT5*-deficient phenotype previously observed in MDA-MB-231 TNBC cells ^22^. shSIRT5 HUVECs also failed to form robust capillary-like networks on Matrigel, with shSIRT5-1 and shSIRT5-2 HUVECs exhibiting reductions in total tube-like segment length of ∼29% (*p <* 0.01) and ∼51% (*p <* 0.0001), respectively, compared with shSC HUVECs. (**Fig 2F-G**). Consistent with these findings with genetic perturbation, pharmacological inhibition of SIRT5 with DK1-04e similarly suppressed proliferation, migration, and tube formation (**Fig. S3A-C**). Together, these results show that SIRT5 is important for ECs to maintain their angiogenic functions.

**Figure 2.**
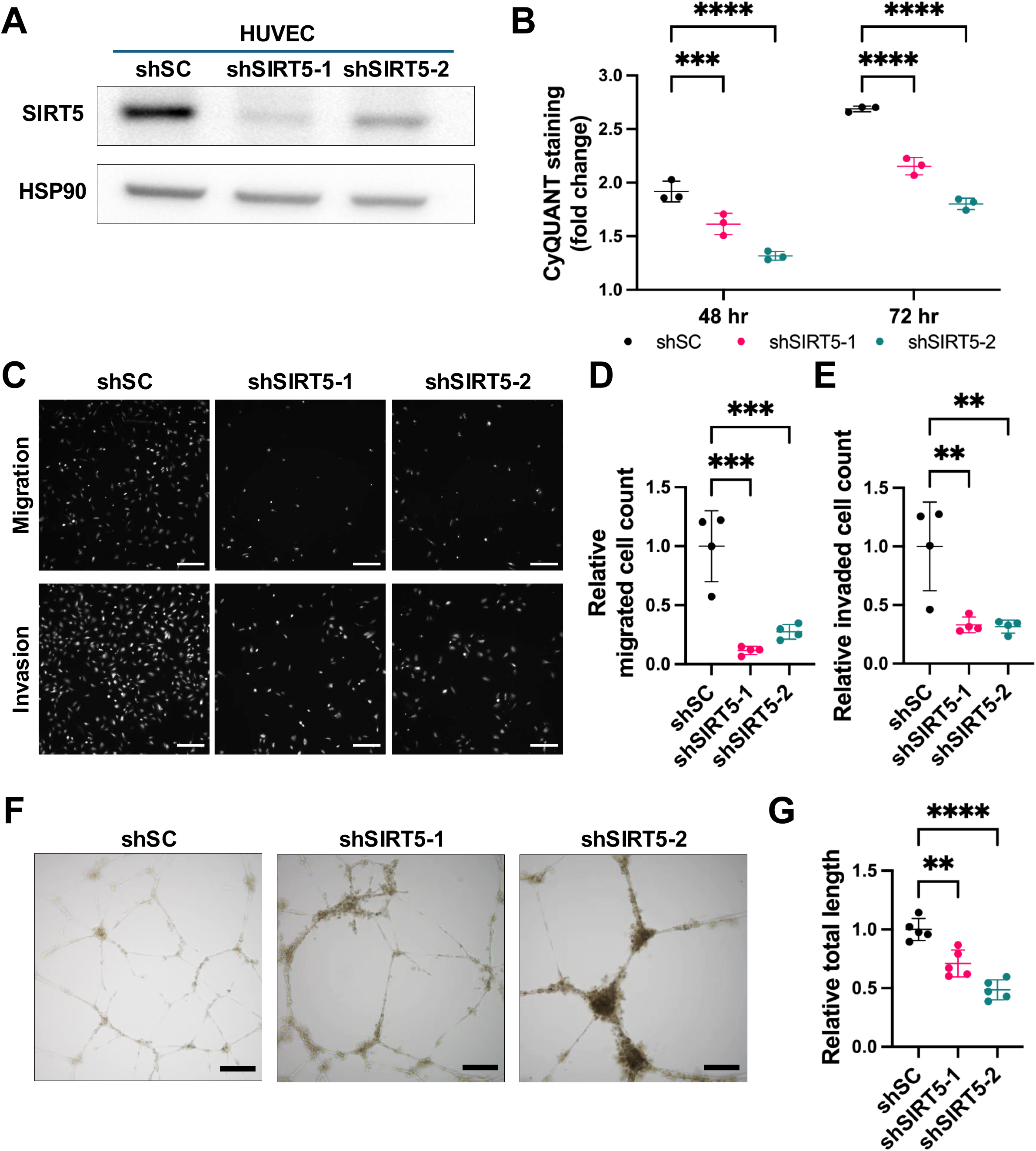
SIRT5 supports endothelial cell angiogenic function *in vitro.* HUVECs were transduced with lentiviral particles carrying two independent shRNA constructs targeting SIRT5 (shSIRT5) or a non-targeting scrambled control (shSC). **(A)** Immunoblot for SIRT5 and HSP90 as a loading control. **(B)** Measurement of viable cell accumulation based on CyQUANT fluorescence intensity at 48 and 72 hours after seeding. Values were normalized to values at 24 hours in each group, respectively (\*\*\*\**p* < 0.0001, two-way ANOVA, n = 3 per group). **(C)** Representative images of migrated and invaded cells at 24 hours post-seeding. Scale bar: 200 µm. **(D-E)** Quantification of relative migrated and invaded cell count in samples from panel (C) (\*\**p <* 0.005, \*\*\**p <* 0.001, one-way ANOVA, n = 4 per group). **(F)** Representative images of the tubular structure formed at 24 hours post-seeding. Scale bar: 200 µm. **(G)** Quantification of relative total segment length of the tubular structure formed in samples from panel (F) (\*\**p <* 0.005, \*\*\*\**p <* 0.001, one-way ANOVA, n = 5 per group).

### SIRT5 inhibition disrupts the ability of EC to maintain vessel barrier integrity and form sprouts in an in vitro three-dimensional microfluidic culture system

To better replicate the physiological microenvironment of blood vessels, which are inherently three-dimensional (3D) structures, we utilized a microfluidic-based 3D culture system ^25,26^ (**Fig. 3A**). The vessel-on-chip device includes two parallel cylindrical microchannels with a collagen hydrogel chamber in a PDMS shell. This platform enables the formation of lumenized, tube-like endothelial structures and incorporates collagen to replicates the extracellular matrix (ECM). Controlled perfusion in this system mimics blood flow and shear stress within the vascular lumen, thereby providing a more physiologically relevant model for studying EC. A representative depth-coded 3D image and corresponding video (IF stained; green: VE-cadherin; blue: DAPI) illustrate the overall architecture of vessels formed in this system (**Fig. 3A, Supplemental video 1**). The structural integrity of vessels formed by shSC and shSIRT5 HUVECs was visualized by VE-cadherin staining and assessed by capturing z-stack fluorescence images of IF-stained vessels 96 hours after seeding and processing them into maximum projections to generate 2D images (**Fig. 3B**). Both shSIRT5-1 and shSIRT5-2 HUVECs formed vessels with prominent cell-free gaps as compared to those formed by shSC HUVECs, which showed almost no gaps **(Fig. 3C)**. These changes indicate a less intact and potentially leakier vessel structure. Although equal numbers of cells were seeded, cell number in shSIRT5 vessels was ∼50-52% lower compared with shSC (*p <* 0.001; **Fig. 3D**). The impact of structural changes in shSIRT5 vessels on vessel function was further investigated by permeability assay as an indicator of barrier integrity. Vessels formed by shSIRT5-1 and shSIRT5-2 HUVECs exhibited increased diffusive permeability by ∼3.6-fold (*p <* 0.001) and ∼3.3-fold (*p <* 0.01), respectively, at t_40_ (**Fig. 3F**), demonstrating that the structural integrity of vessels was impaired upon *SIRT5* loss. Interestingly, these defects were progressive; at 24 hours, barrier integrity of shSIRT5-1 HUVECs did not show significant defects (*p* = 0.2715; **Fig. S4A-B**) despite an early reduction in cell number compared with shSC HUVECs (*p <* 0.05; **Fig. S4C**), suggesting that SIRT5 was particularly critical for the maintenance of mature vessels and that the reduced cell number of shSIRT5 vessels collected at 96 hours was not caused by impaired proliferation alone, possibly due to cell death or changes in cell size. Pharmaceutical inhibition with DK1-04e recapitulated these phenotypes. Vessels formed by DK1-04e treated HUVECs showed a ∼29.6-fold increase in intercellular gap area (*p <* 0.001, **Fig. S5A-B**) and a ∼31% decrease in cell count after 96 hours compared with the vehicle group (*p <* 0.0001, **Fig. S5C**). DK1-04e treated HUVECs increased diffusive permeability by approximately 2.84-fold (*p <* 0.0001) at t_30_ (**Fig. S5D-F**) compared with the vehicle group, validating the results of genetic *SIRT5* inhibition by shRNA (**Fig. 3F**). These findings suggest that SIRT5 loss broadly compromises EC function, impairing both the initial establishment of vessel structures and their maintenance over time.

**Figure 3.**
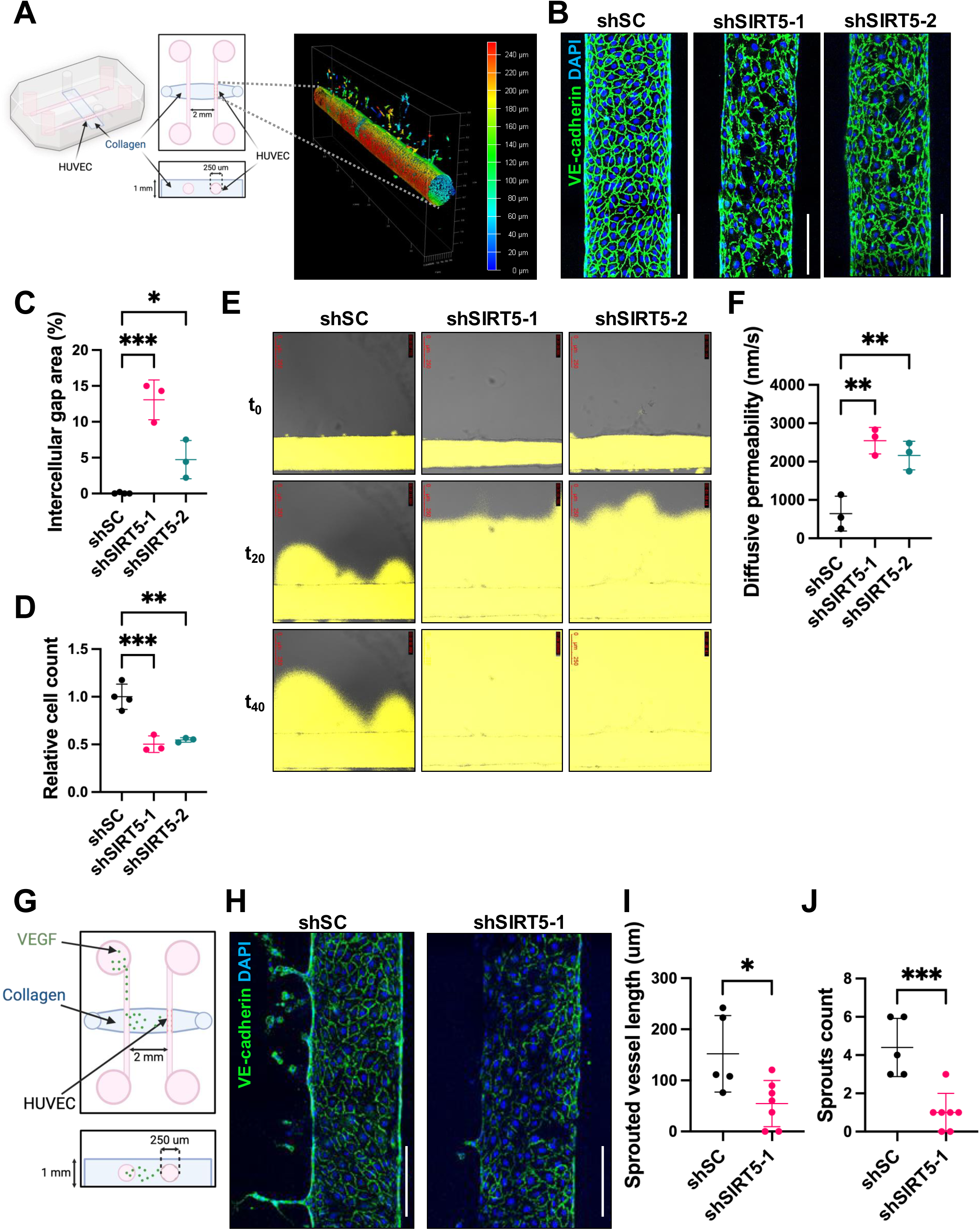
*SIRT5* inhibition disrupts endothelial barrier integrity and sprouting in an *in vitro* 3D microfluidic culture system. shSC and shSIRT5 HUVECs were cultured in the vessel-on-chip model for 96 hours. **(A)** Schematic of the vessel-on-chip 3D *in vitro* culture system. **(B)** Representative images of IF-stained vessels made by HUVECs. Z-stack images of half of each vessel were taken and projected into a single image. Scale bar: 200 µm. **(C-D)** Quantification of the intercellular gap percentage area and relative number of cells in the vessels from panel (B) (\**p <* 0.05, (\*\**p <* 0.01, \*\*\**p <* 0.001, one-way ANOVA, n = 3-4 per group). **(E)** Representative images of permeability assay with fluorescence-labeled dextran at t_0_, t_2_, and t_40_ with vessels. **(F)** Quantification of the permeability in vessels from panel (E) (\*\**p <* 0.005, \*\*\**p <* 0.001, one-way ANOVA, n = 3 per group). **(G)** Design of the sprouting assay with “vessel-on-chip” model. **(H)** Representative images of IF-stained vessels from the vessel sprouting assay. Z-stack images of half of each vessel were taken and projected into a single image. Scale bar: 200 µm. **(I-J)** Quantification of the number and length of sprouts on vessels from panel (H) (\**p <* 0.05, \*\*\**p <* 0.001, one-way ANOVA, n = 5-7 per group). Schematic illustrations were created with BioRender.com.

*SIRT5* KD vessels also failed to initiate robust sprouting. Sprouting was induced by introducing VEGF-A at 20 ng/mL into the chamber without ECs (**Fig. 3G**). Compared with vessels with shSC HUVECs, those with shSIRT5-1 HUVECs exhibited ∼77% (*p <* 0.001) and ∼64% (*p <* 0.05) decreases in the number and length of sprouted vessels, respectively (**Fig. 3H-J**). These data demonstrate that SIRT5 is important for both the structural maintenance and angiogenic switch of 3D vessels.

### SIRT5 regulates mitochondrial respiration and ROS homeostasis in endothelial cells

Since SIRT5 localizes mainly to mitochondria and regulates various metabolic and redox-related proteins ^18^, mitochondrial function in shSC and shSIRT5 HUVECs was next assessed using the Seahorse XFe Mito Stress test (**Fig. 4A**). This test assesses mitochondrial respiratory function in live cells by measuring oxygen consumption in response to sequential perturbations of the electron transport chain. Basal oxygen consumption rate (OCR) was reduced by ∼26% in shSIRT5-1 HUVECs (*p <* 0.001), while shSIRT5-2 HUVECs showed a slight reduction of 2% (*p =* 0.8819, **Fig. 4B**). However, FCCP-stimulated maximal OCR was significantly reduced in both KD lines (shSIRT5-1: ∼64% decrease, *p <* 0.0001; shSIRT5-2: ∼17% decrease, *p <* 0.01). Consequently, spare respiratory capacity was markedly diminished in shSIRT5-1 (∼88% decrease, *p <* 0.0001) and shSIRT5-2 (∼27% decrease, *p <* 0.05) cells (**Fig. 4B**). The degree of these respiratory defects correlated with the KD efficiency of the respective shRNA constructs (**Fig. 2A**). These findings were supported by assessing respiration in parental HUVECs treated with DK1-04e (**Fig. 4C**). Compared with DMSO-treated controls, cells treated with 37.5 µM DK1-04e showed significantly reduced basal OCR (∼52%, *p <* 0.0001), maximal OCR (∼72%, *p <* 0.0001), and spare capacity (∼93%, *p <* 0.001) (**Fig. 4D**). A lower concentration (25 µM DK1-04e) also significantly reduced maximal OCR (∼34%, *p <* 0.01) and spare capacity (∼56%, *p <* 0.01) and caused ∼14% reduction in basal OCR (*p =* 0.1896). These data demonstrate that SIRT5 inhibition impairs mitochondrial respiration in a dose-dependent manner.

**Figure 4.**
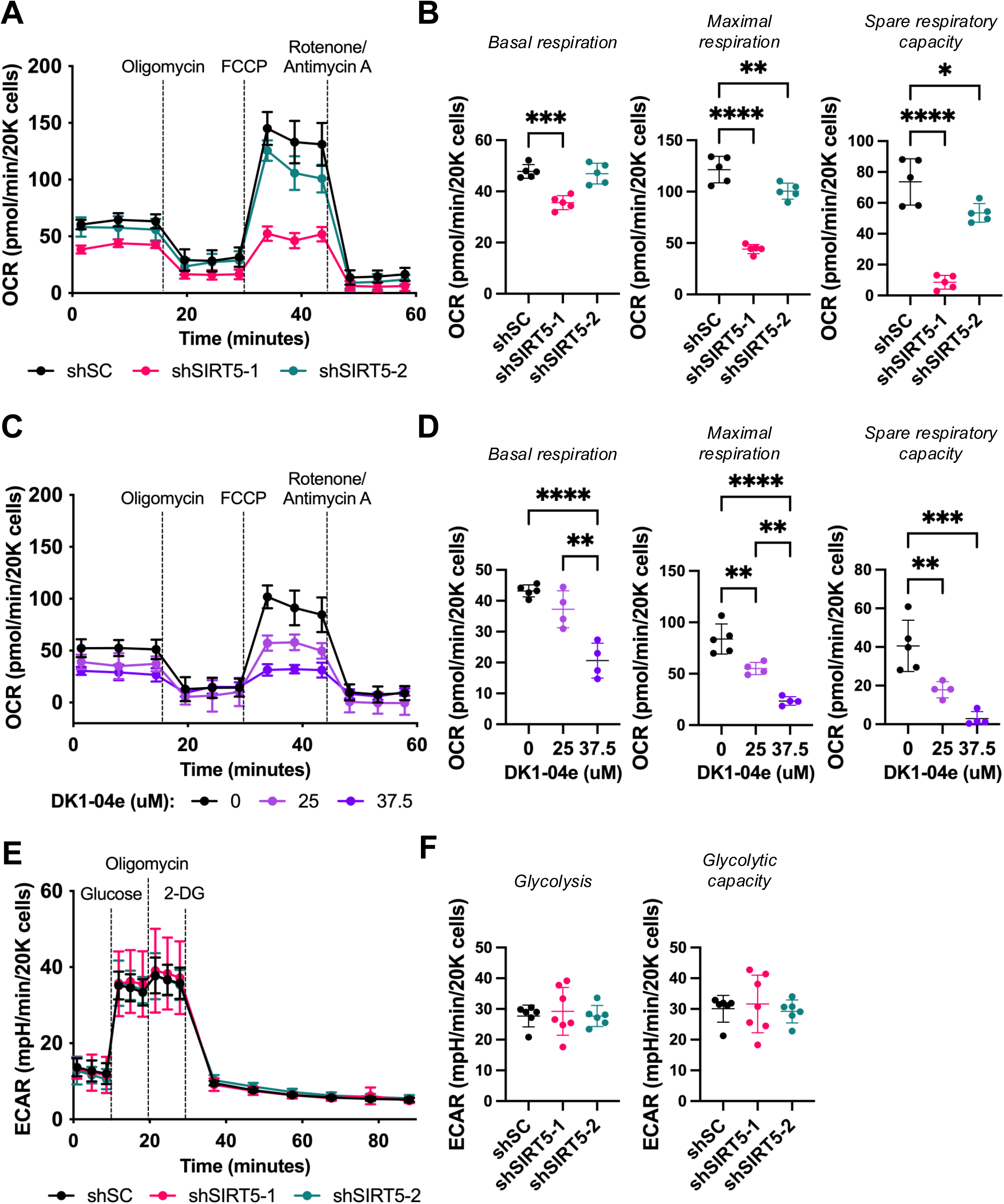
SIRT5 regulates mitochondrial respiration but not glycolytic flux in EC. Mitochondrial function in control and SIRT5-inhibited HUVECs was assessed using the Mito Stress test and flow cytometry. **(A)** Oxygen consumption rate (OCR) results from the Mito Stress test with shSC and shSIRT5 HUVECs. **(B)** Quantification of basal respiration, maximal respiration, and spare respiratory capacity (\**p* < 0.05, \*\**p* < 0.01, \*\*\**p* < 0.001, \*\*\*\**p* < 0.0001, one-way ANOVA, n = 5 per group). **(C)** OCR results from Mito Stress test with DK1-04e and DMSO-treated HUVECs. **(D)** Quantification of basal respiration, maximal respiration, and spare respiratory capacity (\*\**P* value < 0.01, \*\*\**P* value < 0.001, \*\*\*\**P* value < 0.0001, one-way ANOVA, n = 4-5 per group). **(E)** Extracellular acidification rate (ECAR) results from the Glycolytic Stress test with shSC and shSIRT5 HUVECs. **(F)** Quantification of glycolysis and glycolytic capacity (one-way ANOVA, n = 6-7 per group).

The impact of *SIRT5* inhibition on the metabolic fitness of shSC and shSIRT5 HUVECs was further investigated via conducting Seahorse XF Glycolytic Stress Test (**Fig. 4E**). Across all measured parameters, including glycolysis and glycolytic capacity, no significant differences were observed among shSC and the shSIRT5 HUVECs (**Fig. 4F**). Thus, in contrast to mitochondrial respiration, *SIRT5* inhibition does not measurably affect glycolytic function in HUVECs.

Given that mitochondrial respiratory defects (**Fig. 4A-B**) can cause mtROS accumulation and that SIRT5 is a known regulator of redox pathways ^18,33^, we next assessed mtROS levels using MitoSOX and flow cytometry. Under basal conditions, mtROS mean fluorescence intensity (MFI) was significantly elevated in *SIRT5*-deficient HUVECs, increasing ∼1.4-fold in shSIRT5-1 cells and ∼2.5-fold in shSIRT5-2 cells compared with shSC controls (*p* < 0.0001; **Fig. 5A, Fig. S6A**). Treatment with rotenone (0.5 or 1 µM), a complex I inhibitor that increases mitochondrial superoxide, further elevated mtROS in all groups. mtROS levels remained significantly higher in shSIRT5-1 cells (0.5 µM: ∼2.6-fold; 1 µM: ∼1.7-fold) and shSIRT5-2 cells (0.5 µM: ∼2.2-fold; 1 µM: ∼1.6-fold) compared with shSC controls under rotenone treatments (*p* < 0.0001). These data indicate that SIRT5 is required to maintain mitochondrial redox homeostasis under both basal and mitochondrial stress conditions.

**Figure 5.**
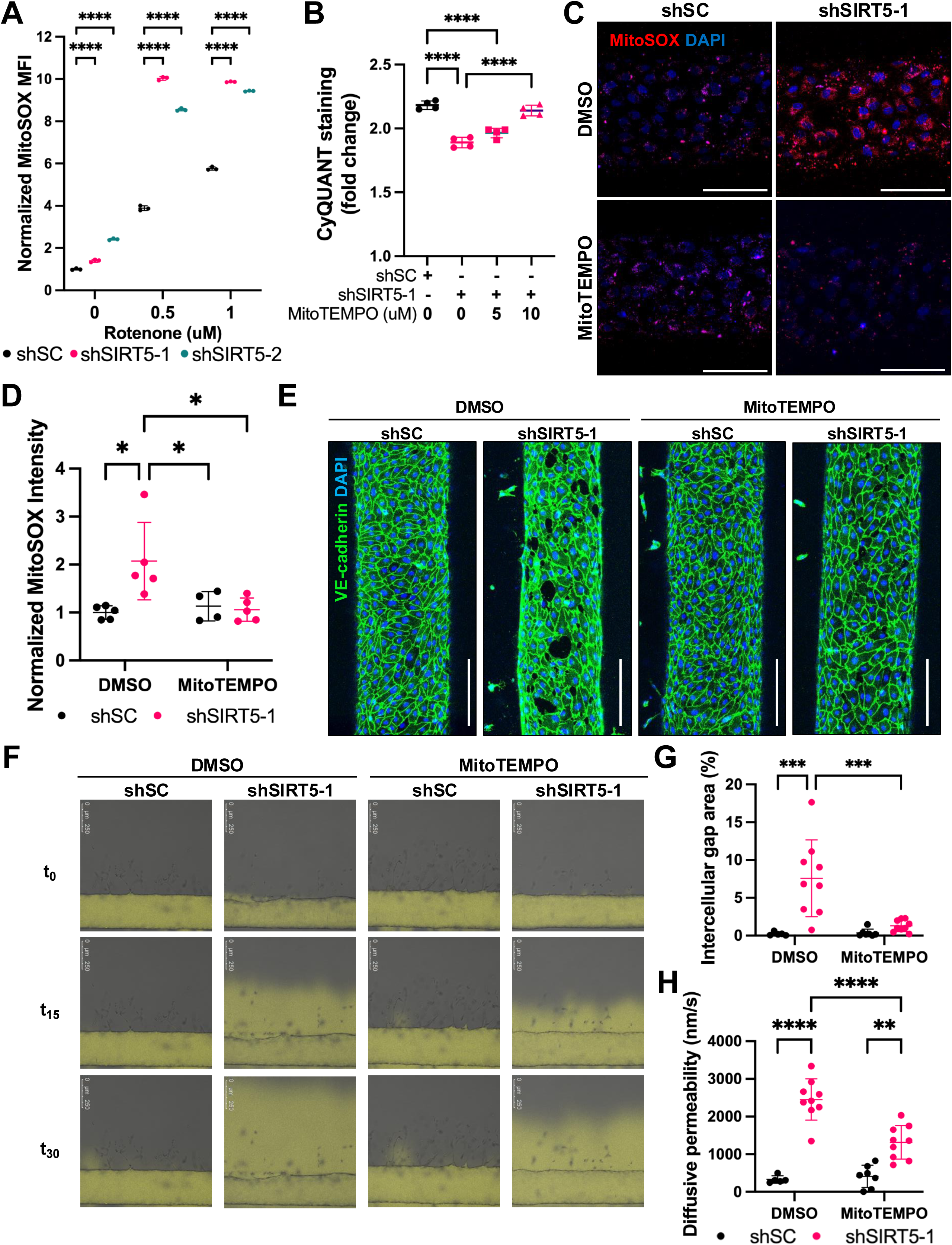
Reducing mtROS rescues the defects of SIRT5-inhibited endothelial cells. shSC and shSIRT5 HUVECs were treated with MitoSOX and/or MitoTEMPO in 2D and the vessel-on-chip culture system. **(A)** Quantification of MitoSOX MFI by flow cytometry (\*\*\*\**p* < 0.0001, one-way ANOVA, n = 3 per group). **(B)** Measurement of viable cell accumulation based on CyQUANT fluorescence intensity at 48 hours after seeding. Values were normalized to the value at 24 hours in each group, respectively (\**p* < 0.05, \*\*\*\**p* < 0.0001, one-way ANOVA, n = 6 per group). **(C)** Representative live-cell images of MitoSOX-stained HUVECs. Z-stack images of half of each vessel were taken and projected into a single image. Scale bar: 200 µm. **(D)** Measurement of MitoSOX Intensity per cell from samples shown in panel (C). (\**p <* 0.05, two-way ANOVA, n = 4-5 devices per group). **(E)** Representative images of IF-stained vessels made by HUVECs. Z-stack images of half of each vessel were taken and projected into a single image. Scale bar: 100 µm. **(F)** Representative images of permeability assay with fluorescence-labeled dextran at t_0_, t_15_, and t_30_ with vessels. **(G)** Quantification of the intercellular percentage gap area in vessels from panel (E) (\*\*\**p <* 0.001, two-way ANOVA, n = 5-8 devices per group). **(H)** Quantification of the permeability in vessels from panel (F) (\**p <* 0.05, \*\**p <* 0.005, **** *p <* 0.001, two-way ANOVA, n = 5-8 per group).

### Defects in SIRT5-inhibited endothelial cells can be rescued by suppressing mtROS levels

To determine whether the observed vascular dysfunction was driven by oxidative stress, we treated *SIRT5* KD HUVECs with the mitochondria-targeted antioxidant MitoTEMPO ^34,35^. Consistent with earlier proliferation assay results, shSIRT5-1 HUVECs treated with the vehicle DMSO showed a ∼25% decrease in proliferation after 48 hours, as compared with shSC cells (**Fig. 5B**, *p <* 0.0001), (**Fig. 2B**). Treatment with 10 µM MitoTEMPO almost fully restored the proliferation capacity of shSIRT5-1 HUVECs, with only a ∼4% decrease in proliferation compared with the vehicle-treated group (**Fig. 5B**, *p =* 0.9522). Treatment with a lower dose of 5 µM MitoTEMPO had an intermediate effect, with the decrease in proliferation in shSIRT5-1 HUVECs diminished to ∼18% of with the vehicle-treated group (**Fig. 5B**, *p <* 0.0001). These results indicate that elevated mtROS contributes to the proliferation impairment in shSIRT5 HUVECs.

The impact of mtROS accumulation in *SIRT5*-deficient ECs was also assessed by adding MitoTEMPO treatment to the vessel-on-chip system (**Fig. 3A**). The mtROS level was first assessed by analyzing live-cell MitoSOX staining via fluorescence imaging (**Fig. 5C)** for later validation of the correlation between mtROS levels and phenotypic changes. The vehicle-treated shSIRT5-1 group showed a ∼2.07-fold increase (**Fig. 5D**, *p <* 0.05) in the MFI of MitoSOX per cell compared with the vehicle-treated shSC group, confirming that *SIRT5* inhibition leads to elevated mtROS levels in HUVECs. However, MitoTEMPO-treated shSIRT5-1 ECs showed no significant differences in the MFI of MitoSOX when compared with both vehicle- and MitoTEMPO-treated shSC ECs, showing that 10 µM MitoTEMPO treatment can reduce MitoROS accumulation in shSIRT5-1 HUVECs.

Treatment with MitoTEMPO significantly rescued the functions of *SIRT5*-deficient ECs. Vehicle-treated shSIRT5-1 HUVECs exhibited severe vascular defects compared with shSC controls, including increased intercellular gaps (7.59% vs. 0.22% area; *p <* 0.001, **Fig. 5E&G**), reduced cell number (∼34%; *p <* 0.0001, **Fig. S6B**), and defective barrier function with increased diffusive permeability (∼7.5-fold; *p <* 0.0001, **Fig. 5F, H**). However, MitoTEMPO treatment (10 μM) reduced intercellular gap area by ∼83% (*p <* 0.0001, **Fig. 5E&G**) and permeability by ∼47% (*p <* 0.0001, **Fig. 5G**) of the shSIRT5-1 vessels compared with vehicle-treated ones, while cell number remained largely unaffected (*p =* 0.0805, **Fig. S6B**). The comparison between treated shSIRT5-1 and shSC vessels revealed that MitoTEMPO fully restored structural integrity (gap area, *p =* 0.9145) but only partially rescued permeability (∼3.2-fold elevation vs. shSC) and cell counts, indicating that mtROS is a primary, but not the only driver of SIRT5-dependent endothelial dysfunction.

### Tumor-extrinsic Sirt5 inhibition in mammary carcinomas increases blood vessel leakiness and impairs tumor oxygen perfusion

Tumor progression requires nutrients and oxygen from the circulation. The impact of SIRT5 deficiency on angiogenesis and vascular function *in vivo* was investigated by assessing tumor vasculature density, vessel permeability, and oxygen perfusion within the TME. Oxygen perfusion and vessel permeability of tumor tissue were assessed by treating mice used in the AT-3 transplant experiment (as shown in **Fig 1A-C**) with Hypoxyprobe^36^, a hypoxia indicator, or fluorescence-labeled dextran^37^, a vascular permeability indicator, before tumor collection. IF staining and imaging of the tumor tissue sections revealed that tumors from *Sirt5* KO hosts showed a 62.5% decrease in CD31-positive area (*p <* 0.01) and ∼5.17-fold increase in Hypoxyprobe-positive area (*p <* 0.05) as compared to those from WT hosts (**Fig. 6A-C**). Fluorescence intensity of dextran was also increased ∼1.64-fold in the tumors from *Sirt5* KO hosts as compared to those from WT hosts (*p <* 0.05; **Fig. 6D-E**). Together, these results demonstrate that host SIRT5 deficiency compromises tumor angiogenesis and vascular integrity, leading to reduced oxygen perfusion and increased permeability, likely due to endothelial dysfunction caused by SIRT5 loss in blood endothelial cells, as supported by our *in vitro* findings.

**Figure 6.**
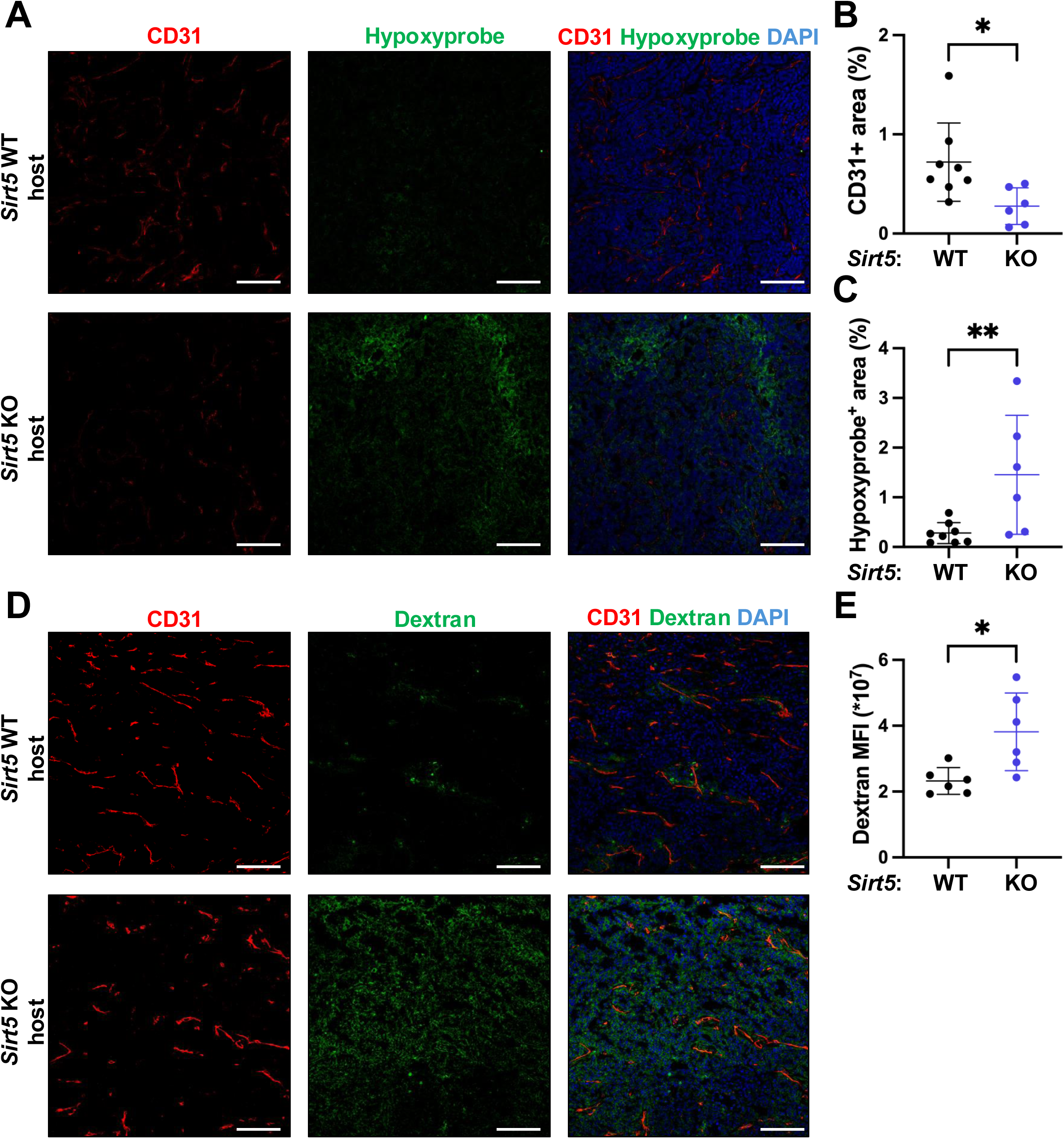
Loss of tumor-extrinsic SIRT5 in mammary carcinoma-bearing mice increases blood vessel leakiness and impairs tumor oxygen perfusion. *Sirt5* WT and KO mice from the syngeneic allograft model with orthotopic AT3 injection (Fig. 1A**-C**) were treated with Hypoxyprobe or fluorescence-labeled dextran before being euthanized. **(A)** Representative IF images of CD31, hypoxyprobe, and DAPI-stained tumors. Scale bar: 100 µm. **(B-C)** Quantification of the mean percentage of CD31-positive area and hypoxyprobe-positive area per field in samples from panel (A) (\**p* < 0.05, \*\**p* < 0.01, unpaired two-tailed Student’s t-test, n = 6-8 per group). **(D)** Representative IF images of CD31 and DAPI-stained tumors with fluorescence-labeled dextran. Scale bar: 100 µm. **(E)** Quantification of the mean of dextran fluorescence intensity per field in samples from panel (D) (\**p* < 0.05, unpaired two-tailed Student’s t-test, n = 6-8 per group).

### SIRT5-deficient ECs fail to support the outgrowth of co-implanted human breast cancer cells in mice

The definitive link of endothelial SIRT5 to BC progression was investigated using a co-injection xenograft mouse model. Orthotopic injection of human MDA-MB-231 TNBC cells with or without HUVECs of varying SIRT5 status into the mammary fat pads of female NSG mice enabled the specific assessment of the effect of endothelial SIRT5 on the growth of engrafted tumors (**Fig. 7A)**. Histopathology revealed severe necrosis and atypia, high mitotic activity, and anisokaryosis in tumors from all 3 groups (**Fig. S7A**). However, tumors without HUVECs progressed slower compared with tumors in both co-implantation groups, while tumors in the MDA-MB-231 + shSIRT5-1 HUVEC group grew significantly slower than those in the MDA-MB-231 + shSC HUVEC group (*p <* 0.001, **Fig. 7B**). Tumors with shSC HUVEC showed ∼2.63-fold (*p <* 0.001) and ∼2.89-fold (*p <* 0.001, **Fig. 7B-C**) greater tumor volume and weight at endpoint compared with the tumors with cancer cells only, demonstrating a pro-tumor role of ECs during tumor progression. However, tumors with shSIRT5-1 HUVECs showed a ∼44% decrease (*p* = 0.1385) and a ∼50% decrease (*p <* 0.01, **Fig. 7B-C**) in tumor volume and weight compared with tumors with shSC HUVECs. This shows that *SIRT5* KD HUVECs are impaired for the ability to support tumor growth when co-injected with MDA-MB-231 cells.

**Figure 7.**
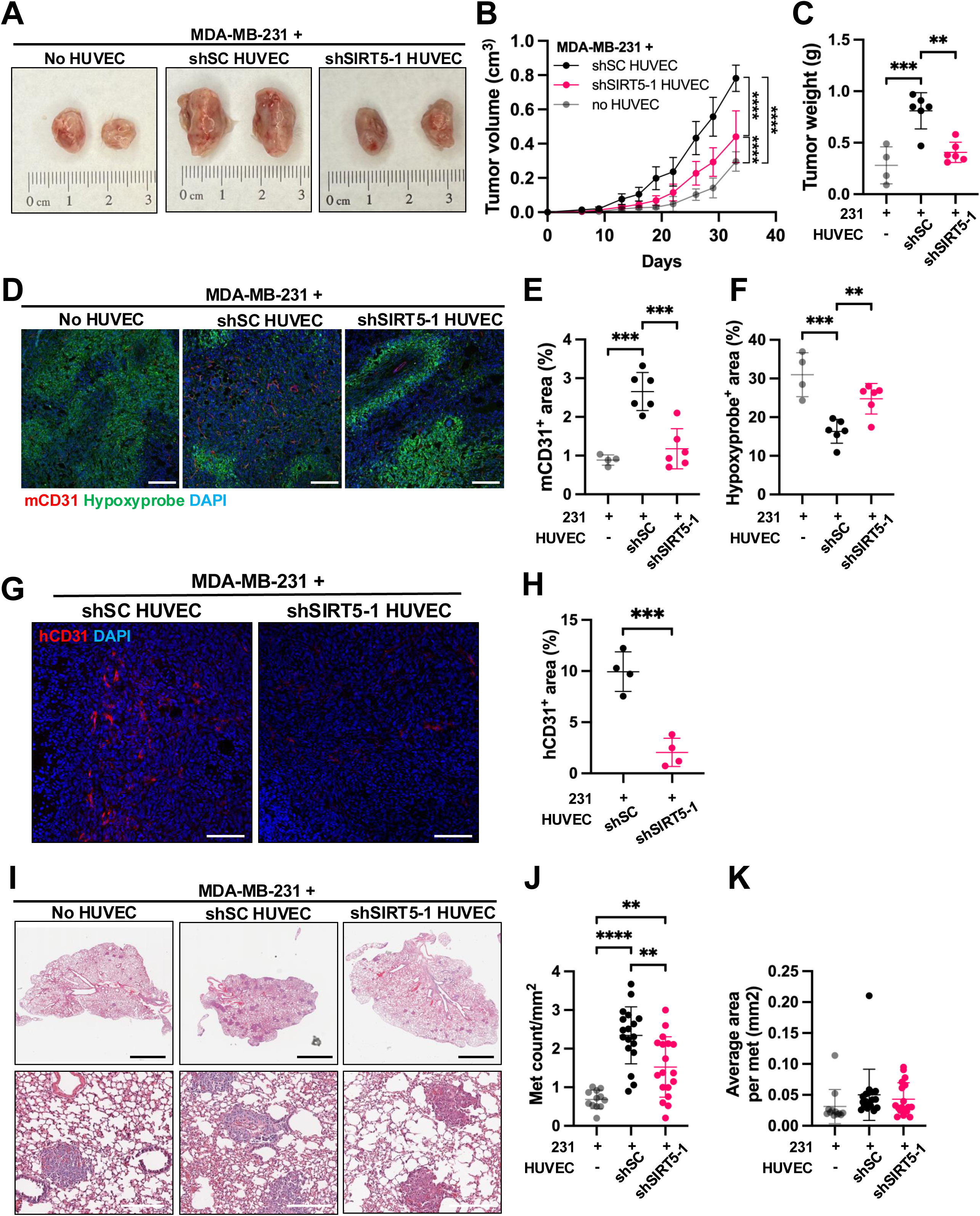
*SIRT5*-deficient ECs fail to support the outgrowth of co-implanted human breast cancer cells in mice. Female NSG mice were orthotopically injected with MDA-MB-231 (231) cells with or without shSC or shSIRT5 HUVECs into the mammary fat pads and treated with Hypoxyprobe before being euthanized. **(A)** Representative images of the tumors in NSG mice collected 35 days post-injection. **(B-C)** Tumor growth curves and weight at endpoint of tumors from panel (A) (\*\**p* < 0.01, \*\*\**p* < 0.001, \*\*\*\**p* < 0.0001, two-way ANOVA, n = 4-6 per group). **(D)** Representative IF images of mCD31, hypoxyprobe, and DAPI-stained tumor sections. Scale bar (white): 200 µm. **(E-F)** Quantification of the mean percentage of CD31-positive area and hypoxyprobe-positive area per field in samples from panel (D) (\*\**p* < 0.01, \*\*\**p* < 0.001, one-way ANOVA, n = 4-6 per group). **(G)** Representative IF images of human CD31 (hCD31) and DAPI-stained tumors collected 18 days post-injection. Scale bar: 100 µm. **(H)** Quantification of the mean percentage of hCD31-positive area per field in samples from panel (G) (\*\*\**p <* 0.001, unpaired two-tailed Student’s t-test, n = 4 per group). **(I)** Representative images of the H&E lung sections collected 35 days post-injection. Scale bar (black): 2 mm. Scale bar (white): 200 µm. **(J-K)** Quantification of met count per mm^2^ lung tissue and average area per met in samples from panel (I) (\*\**p* < 0.01, \*\*\*\**p* < 0.0001, one-way ANOVA, n = 12-18 per group)

To assess the impact of endothelial SIRT5 on vascular function within the TME, we examined tumor vascular density and oxygen perfusion using IF staining for mouse-specific CD31 (mCD31) and Hypoxyprobe (**Fig. 7D)**. Compared with tumors without HUVECs, tumors co-injected with shSC HUVECs showed a ∼3-fold increase (*p <* 0.001, **Fig. 7E**) in mCD31-positive area and a ∼47% reduction in Hypoxyprobe-positive area (*p <* 0.001, **Fig. 7F**), indicating increased angiogenesis and improved oxygenation. In contrast, tumors containing shSIRT5-1 HUVECs showed only a ∼1.3-fold increase in mCD31-positive area (*p =* 0.5842, **Fig. 7E**) and a ∼20% decrease in Hypoxyprobe-positive area (*p =* 0.0869, **Fig. 7F**) relative to MDA-MB-231-only controls. This corresponds to an ∼84% reduction in angiogenic capacity (*p <* 0.001, **Fig. 7E**) and a ∼57% decrease in oxygenation potential (*p <* 0.01, **Fig. 7F**) for shSIRT5-1 HUVECs compared with shSC controls, showing that endothelial SIRT5 was important for inducing more robust tumor angiogenesis and relieving hypoxia. To directly assess the status of the co-implanted HUVECs, we stained the tumors with a human-specific CD31 (hCD31) antibody. However, the hCD31 signal was undetectable in tumors collected at the 35-day endpoint (**Fig. S7B**), possibly reflecting cell death occurring over the prolonged course of the experiment. Hypothesizing that the pro-tumorigenic effects of shSC HUVECs occurred during the earlier stages of tumor growth, we repeated the co-injection experiment with tumor collection at 18 days post-injection (**Fig. S7C**). At this earlier time point, tumor growth and weight were slightly increased in the shSC HUVEC group compared with the no HUVEC and shSIRT5 HUVEC groups (**Fig. S7D-E**). Importantly, HUVECs were observed in tumors from the shSC HUVEC group, as confirmed by hCD31 IF staining. However, hCD31 staining showed a ∼79% reduction in hCD31-positive area in tumors containing shSIRT5-1 HUVECs compared with those with shSC HUVECs (*p <* 0.001, **Fig. 7G-H**). These findings show that *SIRT5*-deficient HUVECs lose their ability to expand or persist within the TME, and that the abundance of HUVECs present during early tumor establishment correlates with subsequent tumor growth and weight at a later time point.

In addition to the primary tumors, we assessed the metastatic potential of MDA-MB-231 cells injected with or without HUVECs of varying SIRT5 status. While shSC HUVECs markedly increased lung metastasis counts (∼3.4-fold vs. no HUVECs, *p <* 0.0001), this effect was significantly reduced in the shSIRT5-1 HUVEC group (∼2.2-fold vs. no HUVECs, *p <* 0.01), showing a ∼34% reduction in metastatic burden compared with the shSC HUVEC group (*p <* 0.01, **Fig. 7I-J**). The average size of individual metastatic lesions did not differ significantly between groups (**Fig. 7K**), suggesting that HUVECs primarily influence the initial dissemination of cancer cells into the circulation, as expected for a model based on EC co-injection at the primary tumor site, rather than their subsequent colonization or proliferation in the lung. These results indicate that loss of endothelial SIRT5 inhibits the ability of ECs to promote both primary tumor progression and metastatic dissemination.

## DISCUSSION

This study establishes the mitochondrial NAD^+^-dependent lysine deacylase SIRT5 as an important regulator of EC metabolic fitness and tumor angiogenesis. We demonstrate that SIRT5 loss in endothelium significantly impairs mammary tumor outgrowth, reduces tumor vascular density, and blunts lung metastatic dissemination. Mechanistically, we show that the absence of SIRT5 impairs the ability of the endothelium to sustain mitochondrial respiration without altering glycolytic flux and leads to abnormal mtROS accumulation. This redox imbalance compromises vascular barrier integrity and sprouting ability, resulting in the formation of structurally defective, leaky vessels that are unable to sustain tumor perfusion. These defects could be rescued by the mitochondria-specific antioxidant MitoTEMPO, indicating that SIRT5-mediated redox homeostasis is an important element of endothelial function. Collectively, our findings show that SIRT5 inhibition can be a novel strategy to dismantle the vascular support system of aggressive BCs, reflecting a targetable metabolic vulnerability of the tumor endothelium.

Despite the important role of SIRT5 in regulating mitochondrial metabolism and redox in ECs, previous characterization of the *Sirt5* KO mouse model revealed no overt defects in developmental or physiological angiogenesis ^25^, suggesting that endothelial SIRT5 is dispensable under non-pathological conditions. This finding is particularly notable when contrasted with the significant impairment of tumor-associated angiogenesis observed in its absence, which strongly suggests a context-dependent requirement for SIRT5. Similarly, DK1-04e was previously shown to be less effective against non-tumorigenic HME1 mammary epithelial cells than MCF7 human breast cancer cells ^22^. Our *in vitro* findings that HUVECs, which are primary cells, rely on SIRT5 under standard culture conditions, while developmental angiogenesis in *Sirt5* KO mice appears grossly normal may reflect the high oxidative stress inherent to *in vitro* systems ^38^. In the context of cancer, the TME imposes severe hypoxia and proliferative stress on ECs during angiogenesis ^4,9,39,40^. Under these intense pathological conditions, the absence of SIRT5 likely sensitizes ECs to metabolic stress, compromising their homeostasis and angiogenic ability. This dependency suggests a promising therapeutic window with limited off-target toxicity for healthy, quiescent vasculature.

Another new finding from this work is the different effects observed for glycolysis and mitochondrial respiration in *SIRT5*-deficient ECs. It is well established that ECs rely primarily on glycolysis for ATP generation—a “Warburg-like” metabolism that supports rapid proliferation and allows ECs to function in hypoxic environments ^10,12,41^. While previous studies show SIRT5 positively regulates glycolysis ^42,43^, SIRT5 loss did not impair glycolytic function in HUVECs. Instead, we observed a selective reduction in mitochondrial respiration. While this diminished respiratory capacity likely restricts overall endothelial metabolism, our rescue experiments demonstrate that the structural vascular defects are dependent on concurrent mtROS accumulation.

While physiological levels of ROS act as signaling molecules to promote angiogenic sprouting ^15^, excessive ROS causes oxidative damage, inflammation, and junctional breakdown. We demonstrate that *SIRT5*-deficient ECs are unable to maintain endothelial redox balance under stressful conditions, leading to barrier dysfunction and the formation of severe intercellular gaps. This phenotype was strikingly evident in our 3D microfluidic model and was validated *in vivo* by increased vascular permeability and hypoxia in *SIRT5*-deficient tumors. In cancer cells, SIRT5 is known to desuccinylate and activate antioxidant enzymes such as SOD1 and IDH2 to prevent oxidative stress-induced destruction ^22,44^. It is highly likely that a similar axis operates in ECs, with SIRT5 supporting the activity of the mitochondrial antioxidant machinery. By linking SIRT5 deficiency directly to ROS-driven endothelial barrier defects, our study offers a mechanistic explanation for how metabolic stress causes vascular leakiness. This may have broad implications not only for tumor angiogenesis but also for vascular pathologies characterized by oxidative stress and barrier failure where roles for SIRT5 have already been established, including ischemic stroke ^45–47^, sepsis ^48^, and colitis ^49^, as well as others where SIRT5 has yet to be implicated, such as vascular aging ^50,51^.

Our observation that global *Sirt5* knockout hosts fail to support tumor growth highlights the importance of SIRT5 in the TME. The concept of tissue-extrinsic roles for SIRT5 in disease is supported by studies of spontaneous and induced cardiac dysfunction, which is observed in global *Sirt5* knockout mice but not in cardiomyocyte-specific SIRT5 knockout mice ^52,53^. While our *in vitro* experiments confirm a significant EC-intrinsic defect, the *in vivo* phenotype observed in the AT-3 transplant model may reflect a composite effect of SIRT5 loss across multiple cell types. In addition to ECs, *SIRT5*-deficient immune cells or stromal cells may also fail to support mammary tumor progression. However, our finding that co-injection of *SIRT5* KD HUVECs alone was sufficient to significantly reduce the ability of EC to promote tumor growth and metastasis demonstrates that the endothelial contribution is a primary driver of the phenotype. This establishes the endothelium as a key therapeutic target for SIRT5 inhibition, even within a complex multicellular TME. Notably, previous research has demonstrated that ECs can actually inhibit breast cancer cell growth via angiocrine regulation *in vitro* ^54^. This suggests that the pro-tumorigenic effects of endothelial cells observed in our co-injection model are likely caused by their structural effects on tumor vascularization, a function that is critically impaired by SIRT5 perturbation.

Current anti-VEGF therapies often fail because they induce excessive vascular regression, leading to hypoxia that promotes cancer cell aggressiveness and impairs antitumor immunity, or because tumors develop resistance by secreting alternative angiogenic factors. SIRT5 inhibition represents a fundamentally different approach: targeting endothelial metabolic fitness rather than growth factor signaling. We observed that severe SIRT5 perturbation leads to a non-functional, highly permeable vasculature that fails to support tumor expansion. However, prior studies have demonstrated that while high doses of anti-angiogenic drugs suppress angiogenesis, moderate dosing can normalize the tumor vasculature ^55–57^. The concept of “vascular normalization”—pruning of immature, leaky vessels to improve perfusion, immune infiltration, and drug delivery—is a novel therapeutic goal and functional outcome of anti-angiogenic therapy ^58–60^. It is possible that a calibrated pharmacological inhibition of SIRT5 could be evaluated as a potential vascular normalization method. Importantly, because ECs are genetically stable and less prone to mutation than cancer cells, a SIRT5-targeted therapy directed at the endothelium may be less susceptible to the rapid development of drug resistance that plagues therapies targeting tumor cells directly.

To summarize, this work shows SIRT5 as a key regulator of endothelial redox homeostasis. We define a specific metabolic vulnerability in the ECs: a reliance on SIRT5-mediated buffering to mitigate oxidative stress. By identifying SIRT5 as an important metabolic regulator that is dispensable for normal physiology but vital for pathological angiogenesis, we provide a rationale for developing SIRT5 inhibitors as a new class of anti-angiogenic agents. Coupled with prior evidence that SIRT5 supports TNBC cell proliferation and invasiveness^22^ and the minimal side-effects in SIRT5-disrupted mice, targeting SIRT5 could dismantle both the vasculature and the tumor-intrinsic growth machinery of TNBC. This provides a dual-action mechanism to suppress primary tumors and prevent metastatic dissemination and has the potential to be an optimal strategy for novel TNBC treatment.

## ACKNOWLEDGEMENTS

This study was supported by funding from NIH grants R01 CA223534 to **RSW**, T32 ODO011000 to **AMC**, and R01 HL165135 to **EL**. We acknowledge the Center for Animal Resources and Education (CARE) and the Flow Cytometry Facility at Cornell University for help with the study. Dr. Praveen Sethupathy is acknowledged for providing tissue specimens. Andy Chang, Scott Butler, Kelsey Jones, and Evan Zhou are acknowledged for their assistance with experiments and Ravi Dhawan for contributions to preliminary studies.

## AUTHOR CONTRIBUTIONS

A.M.C. and R.S.W. conceptualized the project. A.M.C., I.C., Q.Z., P.T., Z.C.N., J.B., E.L., and R.S.W. developed the methodology. A.M.C., I.C., Q.Z., E.A.B., and I.R.F. performed the investigation. A.M.C. validated the project. A.M.C. and A.M. conducted the formal analysis. A.M.C. curated the data. A.M.C. provided visualization. A.M.C. wrote the original draft of the manuscript; all authors reviewed and edited the final version. A.M.C., S.J., Z.C.N., J.B., H.L., E.L., and R.S.W. acquired resources. E.L. and R.S.W. supervised the work. E.L. and R.S.W. acquired funding.

## REFERENCES

1. Bray Bsc, F., et al. Global cancer statistics 2022: GLOBOCAN estimates of incidence and mortality worldwide for 36 cancers in 185 countries. CA Cancer J. Clin. 74, 229–263 (2024).

2. Zagami, P. & Carey, L. A. Triple negative breast cancer: Pitfalls and progress. npj Breast Cancer 2022 8:1 8, 95- (2022).

3. Ribatti, D. et al. Angiogenesis and Antiangiogenesis in Triple-Negative Breast cancer. Transl. Oncol. 9, 453 (2016).

4. Hanahan, D. & Weinberg, R. A. Hallmarks of cancer: The next generation. Cell 144, 646–674 (2011).

5. Sasich, L. D. & Sukkari, S. R. The US FDAs withdrawal of the breast cancer indication for Avastin (bevacizumab). Saudi Pharm. J. 20, 381–385 (2012).

6. Liu, Z. L., Chen, H. H., Zheng, L. L., Sun, L. P. & Shi, L. Angiogenic signaling pathways and anti-angiogenic therapy for cancer. Signal Transduction and Targeted Therapy 2023 8:1 8, 1–39 (2023).

7. Vimalraj, S. A concise review of VEGF, PDGF, FGF, Notch, angiopoietin, and HGF signalling in tumor angiogenesis with a focus on alternative approaches and future directions. Int. J. Biol. Macromol. 221, 1428–1438 (2022).

8. Garcia, J. et al. Bevacizumab (Avastin®) in cancer treatment: A review of 15 years of clinical experience and future outlook. Cancer Treat. Rev. 86, 102017 (2020).

9. Kane, K., Edwards, D. & Chen, J. The influence of endothelial metabolic reprogramming on the tumor microenvironment. Oncogene 2024 44:2 44, 51–63 (2024).

10. Li, X., Sun, X. & Carmeliet, P. Hallmarks of Endothelial Cell Metabolism in Health and Disease. Cell Metab. 30, 414–433 (2019).

11. García-Caballero, M., Sokol, L., Cuypers, A. & Carmeliet, P. Metabolic Reprogramming in Tumor Endothelial Cells. Int. J. Mol. Sci. 23, (2022).

12. Warburg, O., Wind, F. & Negelein, E. The Metabolism of Tumors in the Body. J. Gen. Physiol. 8, 519–530 (1927).

13. Kluge, M. A., Fetterman, J. L. & Vita, J. A. Mitochondria and Endothelial Function. Circ. Res. 112, 1171 (2013).

14. Connor, K. M. et al. Mitochondrial H2O2 regulates the angiogenic phenotype via PTEN oxidation. J. Biol. Chem. 280, 16916–16924 (2005).

15. Panieri, E. & Santoro, M. M. ROS signaling and redox biology in endothelial cells. Cell. Mol. Life Sci. 72, 3281 (2015).

16. Minjares, M., Wu, W. & Wang, J. M. Oxidative Stress and MicroRNAs in Endothelial Cells under Metabolic Disorders. Cells 12, 1341 (2023).

17. Wang, M. & Lin, H. Understanding the Function of Mammalian Sirtuins and Protein Lysine Acylation. Annu. Rev. Biochem. 90, 245–285 (2021).

18. Fabbrizi, E. et al. Emerging Roles of SIRT5 in Metabolism, Cancer, and SARS-CoV-2 Infection. Cells 12, 852 (2023).

19. Du, J. et al. Sirt5 Is an NAD-Dependent Protein Lysine Demalonylase and Desuccinylase. Science 334, 806 (2011).

20. Giblin, W. et al. The deacylase SIRT5 supports melanoma viability by influencing chromatin dynamics. J. Clin. Invest. 131, e138926 (2021).

21. Yan, D. et al. SIRT5 Is a Druggable Metabolic Vulnerability in Acute Myeloid Leukemia. Blood Cancer Discov. 2, 66–87 (2021).

22. Abril, Y. L. N. et al. Pharmacological and genetic perturbation establish SIRT5 as a promising target in breast cancer. Oncogene 2021 40:9 40, 1644–1658 (2021).

23. Greene, K. S. et al. SIRT5 stabilizes mitochondrial glutaminase and supports breast cancer tumorigenesis. Proc. Natl. Acad. Sci. U. S. A. 116, 26625–26632 (2019).

24. Lagunas-Rangel, F. A. Role of SIRT5 in cancer. Friend or Foe? Biochimie 209, 131–141 (2023).

25. Yu, J. et al. Metabolic Characterization of a Sirt5 deficient mouse model. Scientific Reports 2013 3:1 3, 1–7 (2013).

26. Davie, S. A. et al. Effects of FVB/NJ and C57Bl/6J strain backgrounds on mammary tumor phenotype in inducible nitric oxide synthase deficient mice. Transgenic Res. 16, 193–201 (2007).

27. Stewart, T. J. & Abrams, S. I. Altered Immune Function during Long-Term Host-Tumor Interactions Can Be Modulated to Retard Autochthonous Neoplastic Growth. The Journal of Immunology 179, 2851–2859 (2007).

28. Stewart, T. J. & Abrams, S. I. Altered Immune Function during Long-Term Host-Tumor Interactions Can Be Modulated to Retard Autochthonous Neoplastic Growth. The Journal of Immunology 179, 2851–2859 (2007).

29. Lee, E. et al. A 3D biomimetic model of lymphatics reveals cell–cell junction tightening and lymphedema via a cytokine-induced ROCK2/JAM-A complex. Proc. Natl. Acad. Sci. U. S. A. 120, e2308941120 (2023).

30. Polacheck, W. J., Kutys, M. L., Tefft, J. B. & Chen, C. S. Microfabricated blood vessels for modeling the vascular transport barrier. Nature Protocols 2019 14:5 14, 1425–1454 (2019).

31. Qu, Y. et al. Small molecule oxybutynin rescues proliferative capacity of complex III-defective muscle progenitor cells. Am. J. Physiol. Cell Physiol. 329, C911–C923 (2025).

32. Cheng, C. et al. AQP1: a regulatory factor associated with brown adipose tissue-silencing to combat obesity and metabolic disease. Am. J. Physiol. Endocrinol. Metab. 329, (2025).

33. De Filippis, B. et al. Mitochondrial free radical overproduction due to respiratory chain impairment in the brain of a mouse model of Rett syndrome: protective effect of CNF1. Free Radic. Biol. Med. 83, 167–177 (2015).

34. Wang, X. et al. Scinderin promotes fusion of electron transport chain dysfunctional muscle stem cells with myofibers. *Nat*. Aging 2, 155 (2022).

35. Peng, X. et al. HMOX1-LDHB interaction promotes ferroptosis by inducing mitochondrial dysfunction in foamy macrophages during advanced atherosclerosis. Dev. Cell 60, 1070–1086.e8 (2025).

36. Oza, H. H., Ng, E. & Gilkes, D. M. Staining Hypoxic Areas of Frozen and FFPE Tissue Sections with Hypoxyprobe^TM^. Methods Mol. Biol. 2755, 149–163 (2024).

37. Dreher, M. R. et al. Tumor vascular permeability, accumulation, and penetration of macromolecular drug carriers. J. Natl. Cancer Inst. 98, 335–344 (2006).

38. Halliwell, B. Oxidative stress in cell culture: an under-appreciated problem? FEBS Lett. 540, 3–6 (2003).

39. Lugano, R., Ramachandran, M. & Dimberg, A. Tumor angiogenesis: causes, consequences, challenges and opportunities. Cell. Mol. Life Sci. 77, 1745 (2019).

40. Hanahan, D. & Folkman, J. Patterns and emerging mechanisms of the angiogenic switch during tumorigenesis. Cell 86, 353–364 (1996).

41. De Bock, K., Georgiadou, M. & Carmeliet, P. Role of Endothelial Cell Metabolism in Vessel Sprouting. Cell Metab. 18, 634–647 (2013).

42. Yan, D. et al. SIRT5 Is a Druggable Metabolic Vulnerability in Acute Myeloid Leukemia. Blood Cancer Discov. 2, 266 (2021).

43. He, S., Jia, Q., Zhou, L., Wang, Z. & Li, M. SIRT5 is involved in the proliferation and metastasis of breast cancer by promoting aerobic glycolysis. Pathol. Res. Pract. 235, (2022).

44. Lin, Z. F. et al. SIRT5 desuccinylates and activates SOD1 to eliminate ROS. Biochem. Biophys. Res. Commun. 441, 191–195 (2013).

45. Zhang, L., Lv, T., Hou, P., Jin, Y. & Jia, F. Sirt5-mediated polarization and metabolic reprogramming of macrophage sustain brain function following ischemic stroke. Brain Res. 1857, 149613 (2025).

46. Li, J. et al. SIRT5 Regulates Ferroptosis through the Nrf2/HO-1 Signaling Axis to Participate in Ischemia-Reperfusion Injury in Ischemic Stroke. Neurochemical Research 2024 49:4 49, 998–1007 (2024).

47. Morris-Blanco, K. C. et al. Protein Kinase C Epsilon Promotes Cerebral Ischemic Tolerance Via Modulation of Mitochondrial Sirt5. Scientific Reports 2016 6:1 6, 29790- (2016).

48. Zhang, X. et al. Desuccinylation of TBK1 by SIRT5 regulates inflammatory response of macrophages in sepsis. Cell Rep. 43, 115060 (2024).

49. Wang, F. et al. SIRT5 Desuccinylates and Activates Pyruvate Kinase M2 to Block Macrophage IL-1β Production and to Prevent DSS-Induced Colitis in Mice. Cell Rep. 19, 2331–2344 (2017).

50. Oakley, R. & Tharakan, B. Vascular hyperpermeability and aging. Aging Dis. 5, 114–125 (2014).

51. Ungvari, Z., Tarantini, S., Donato, A. J., Galvan, V. & Csiszar, A. Mechanisms of Vascular Aging. Circ. Res. 123, 849–867 (2018).

52. Sadhukhan, S. et al. Metabolomics-assisted proteomics identifies succinylation and SIRT5 as important regulators of cardiac function. Proc. Natl. Acad. Sci. U. S. A. 113, 4320–4325 (2016).

53. Hershberger, K. A. et al. Ablation of Sirtuin5 in the postnatal mouse heart results in protein succinylation and normal survival in response to chronic pressure overload. J. Biol. Chem. 293, 10630 (2018).

54. Franses, J. W., Baker, A. B., Chitalia, V. C. & Edelman, E. R. Stromal Endothelial Cells Directly Influence Cancer Progression. Sci. Transl. Med. 3, 66ra5 (2011).

55. Cantelmo, A. R. et al. Inhibition of the Glycolytic Activator PFKFB3 in Endothelium Induces Tumor Vessel Normalization, Impairs Metastasis, and Improves Chemotherapy. Cancer Cell 30, 968–985 (2016).

56. Matsumoto, K. et al. Inhibition of glycolytic activator PFKFB3 suppresses tumor growth and induces tumor vessel normalization in hepatocellular carcinoma. Cancer Lett. 500, 29–40 (2021).

57. Conradi, L. C. et al. Tumor vessel disintegration by maximum tolerable PFKFB3 blockade. Angiogenesis 20, 599–613 (2017).

58. Jain, R. K. Normalization of tumor vasculature: an emerging concept in antiangiogenic therapy. Science 307, 58–62 (2005).

59. Jain, R. K. Normalizing tumor vasculature with anti-angiogenic therapy: A new paradigm for combination therapy. Nat. Med. 7, 987–989 (2001).

60. Carmeliet, P. & Jain, R. K. Principles and mechanisms of vessel normalization for cancer and other angiogenic diseases. Nature Reviews Drug Discovery 2011 10:6 10, 417–427 (2011).

