## Supplemental Information for "SIRT5 acts in the tumor microenvironment via endothelial cell metabolism to support breast cancer growth"

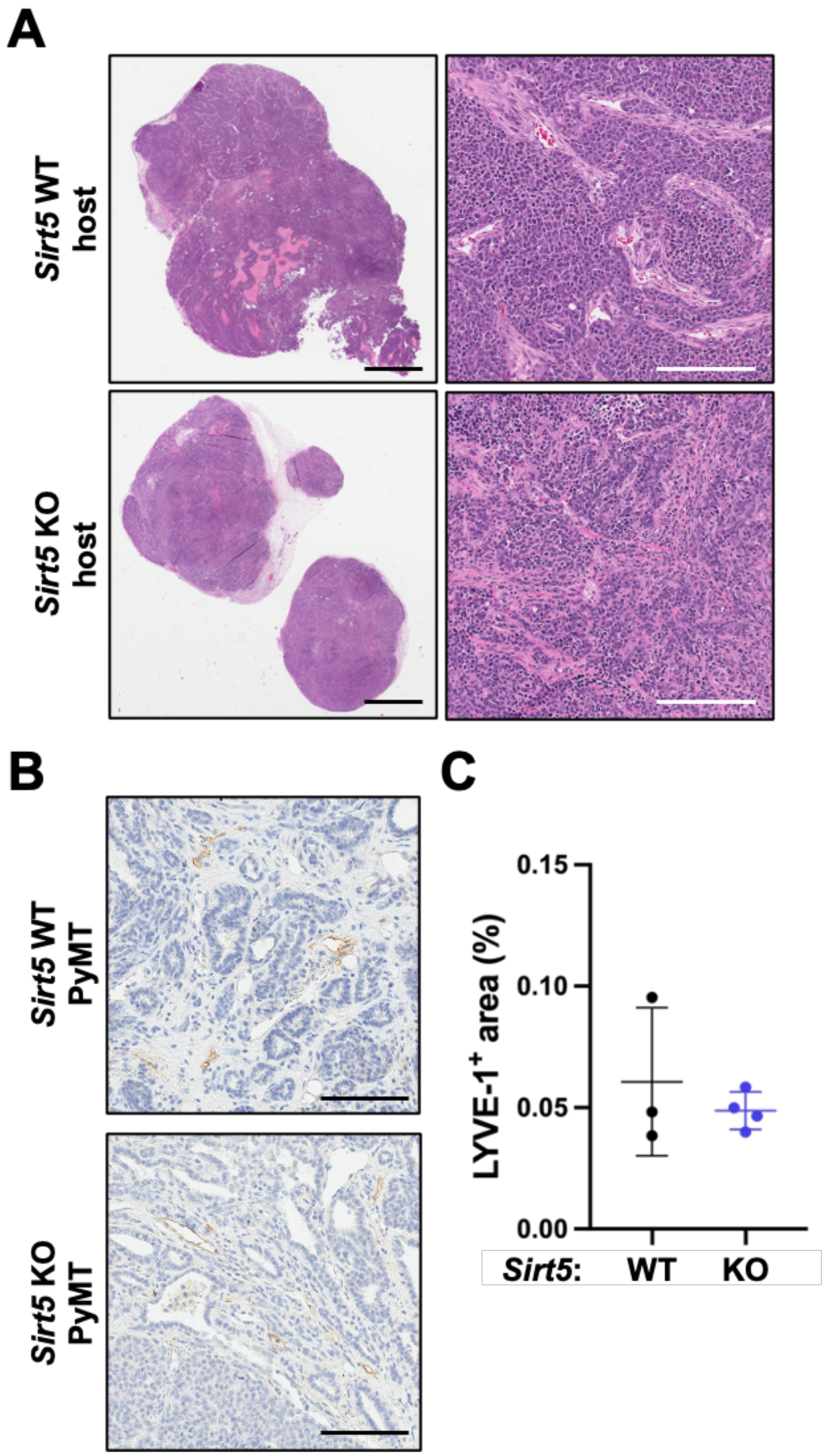


**Figure S1. *Sirt5* inhibition does not significantly alter lymphangiogenesis in mouse mammary carcinomas. (A)** Representative H&E images of tumor sections from syngeneic orthotopic AT3 mammary tumors in *Sirt5* WT and KO mice. Scale bar (black):

2 mm. Scale bar (white): 200 μm. **(B)** Representative LYVE-1 IHC images of mammary tumors from WT and *Sirt5* KO PyMT mice. Scale bar, 100 μm. **(C)** Quantification of Lyve-1-positive area in tumors from panel (B) (unpaired two-tailed Student's t-test).

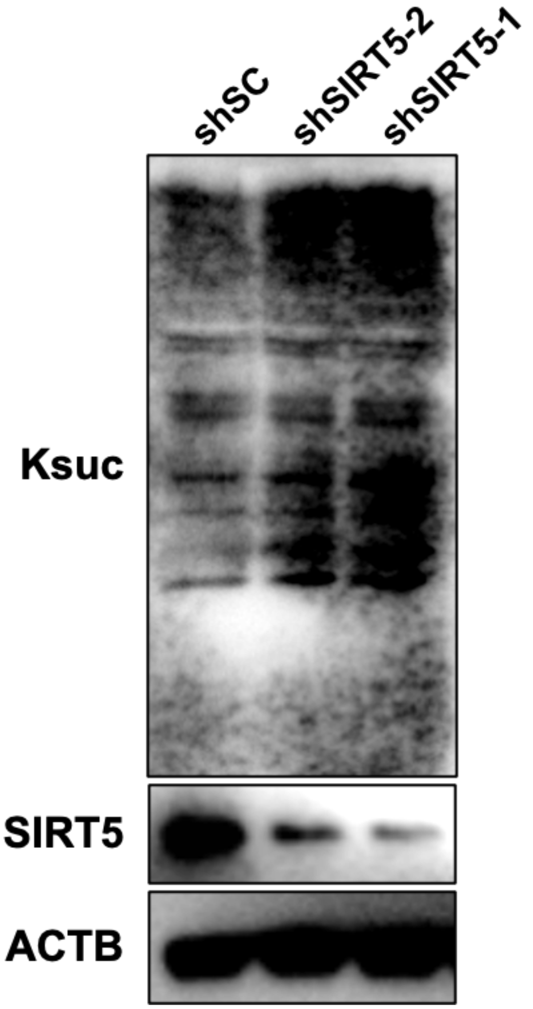


**Figure S2. SIRT5 inhibition leads to elevated lysine succinylation (Ksuc) in endothelial cells.** Protein lysates from shSC and shSIRT5 HUVECs were immunoblotted for lysine succinylation (Ksuc) and SIRT5 to assess the correlation between global Ksuc and SIRT5 levels. ACTB was used as a loading control.

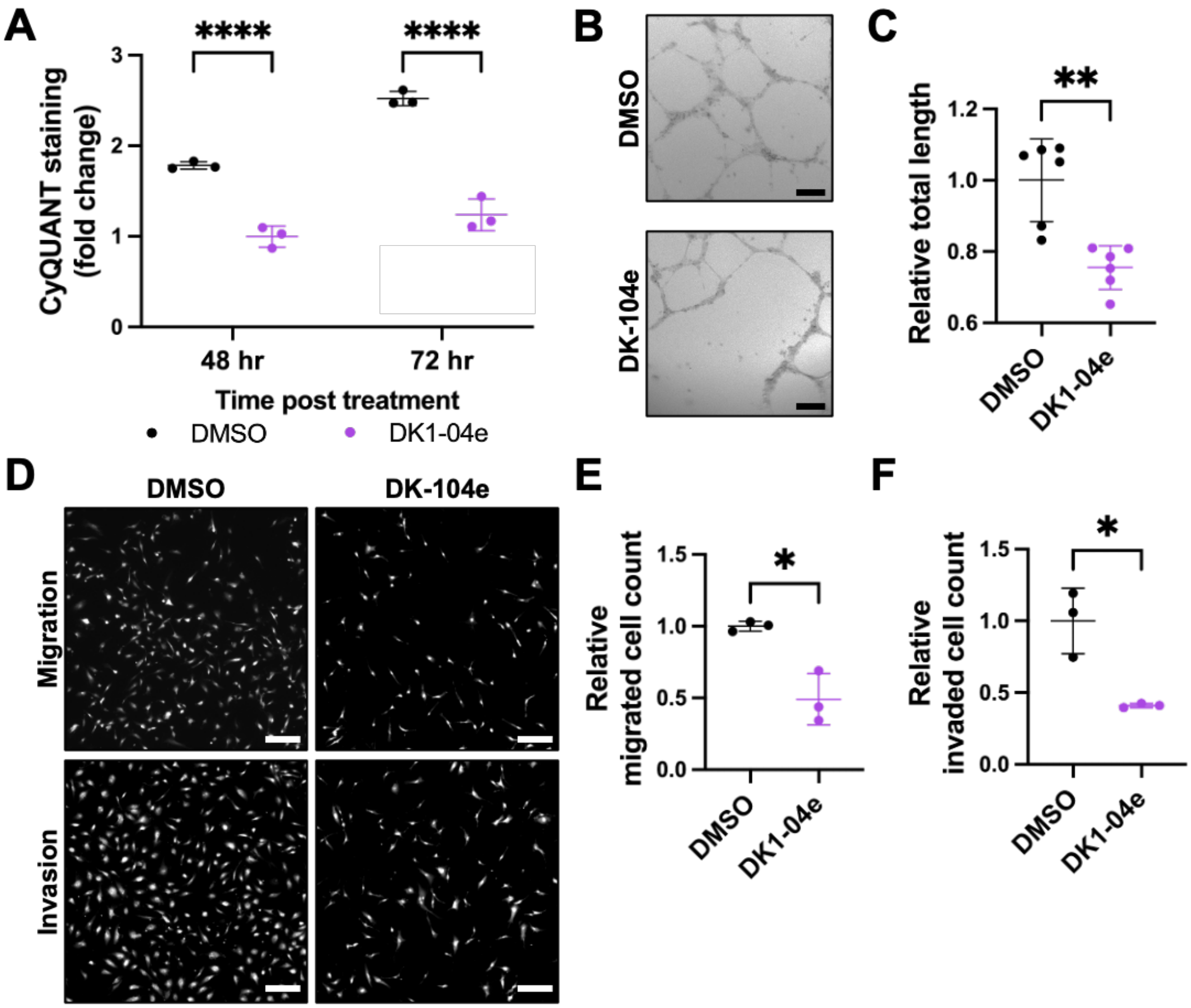


**Figure S3. SIRT5-selective inhibitor DK1-04e reduces angiogenic functions of HUVECs.** HUVECs were treated with DK1-04e or DMSO (vehicle control). **(A)** Measurement of viable cell accumulation based on CyQUANT fluorescence intensity at 48 and 72 hours after seeding. The values were normalized to the value at 24 hours in each group, respectively (*****P* value < 0.0001, two-way ANOVA, n = 3 per group). **(B)** Representative images of the tubular structure formed at 24 hours post-seeding. Scale bar: 200 µm. **(C)** Quantification of relative total segment length of the tubular structure in samples of panel (B) (***P* value < 0.005, one-way ANOVA, n = 5 per group). **(D)** Representative images of migrated and invaded cells stained with PI at 24 hours post-seeding. Scale bar: 200 μm. **(E-F)** Quantification of relative migrated and invaded cell count in samples from panel (D) (**P* value < 0.05, ***P* value < 0.005, one-way ANOVA, n = 4 per group).

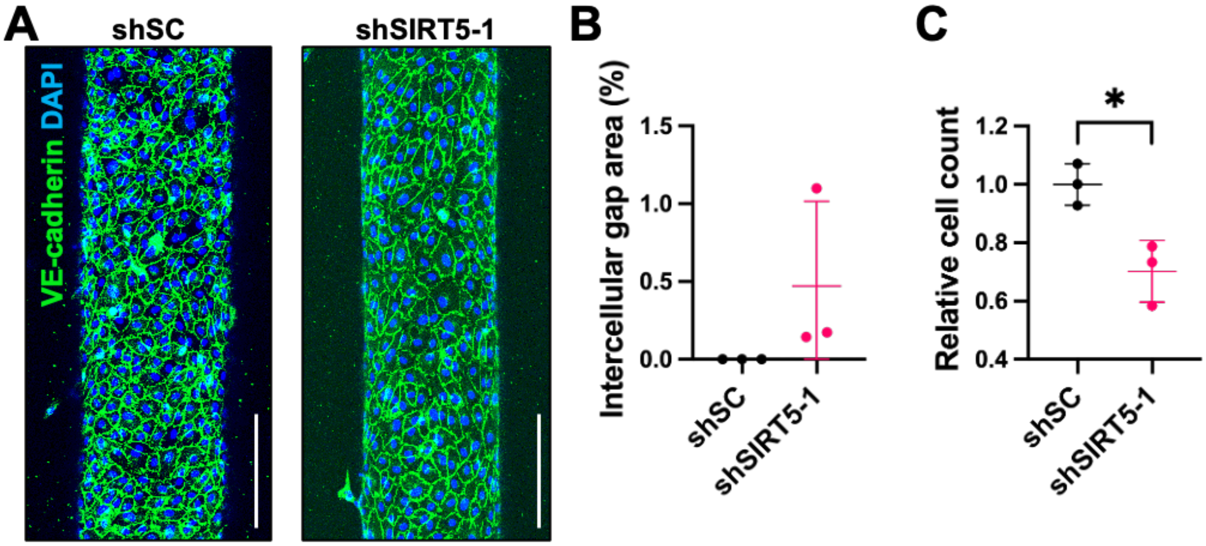


**Figure S4. *SIRT5* loss in ECs causes decreased cell number but not a significant increase in intercellular gap area at the early time point in an *in vitro* 3D microfluidic culture system.** shSC and shSIRT5 HUVECs were cultured in the vessel-on-chip model for 24 hours. **(A)** Representative images of IF-stained vessels made by HUVECs. Z-stack images of half of each vessel were taken and projected into a single image. Scale bar: 200 µm. **(B-C)** Quantification of the intercellular gap percentage area and the relative number of cells in the vessels of panel (A) (**P* value < 0.05, unpaired two-tailed Student's t-test, n = 3 per group).

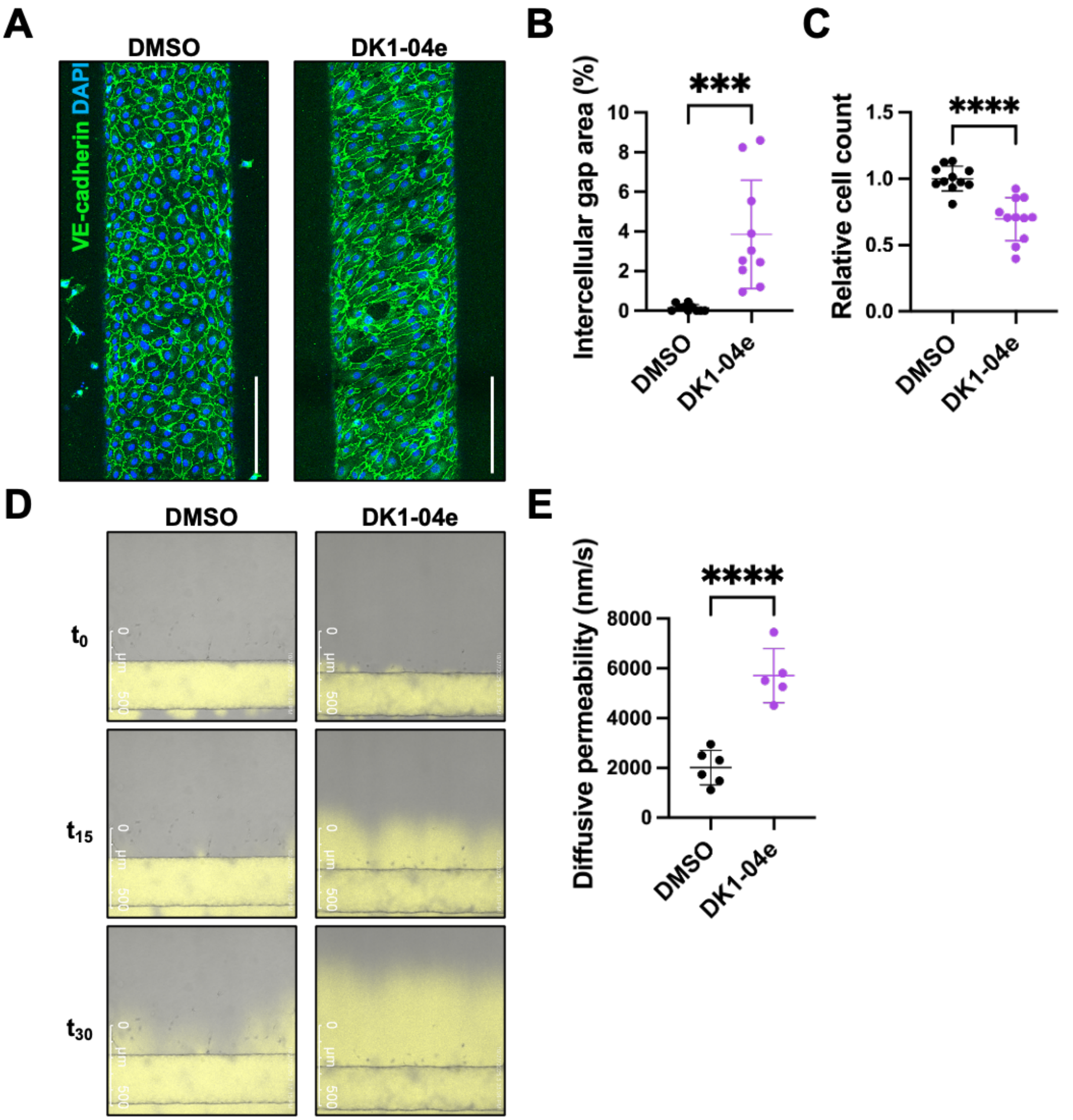


**Figure S5. SIRT5-selective inhibitor DK1-04e disrupts endothelial cell ability to maintain vessel barrier integrity in an *in vitro* 3D microfluidic culture system.** DK1-04e and vehicle (DMSO) treated HUVECs were cultured in the vessel-on-chip model for 96 hours. **(A)** Representative images of IF-stained vessels made by HUVECs. Z-stack images of half of each vessel were taken and projected into a single image. Scale bar:

200 μm. **(B-C)** Quantification of the intercellular gap percentage area and the relative number of cells in the vessels of panel (A) (****P* value < 0.001, unpaired two-tailed Student's t-test, n = 10 per group). **(D)** Representative images of permeability assay with fluorescence-labeled dextran at $t_0$, $t_{15}$, and $t_{30}$ with vessels. **(E)** Quantification of the permeability in vessels of panel (D) (**** *P* value < 0.0001, one-way ANOVA, n = 5-6 per group).

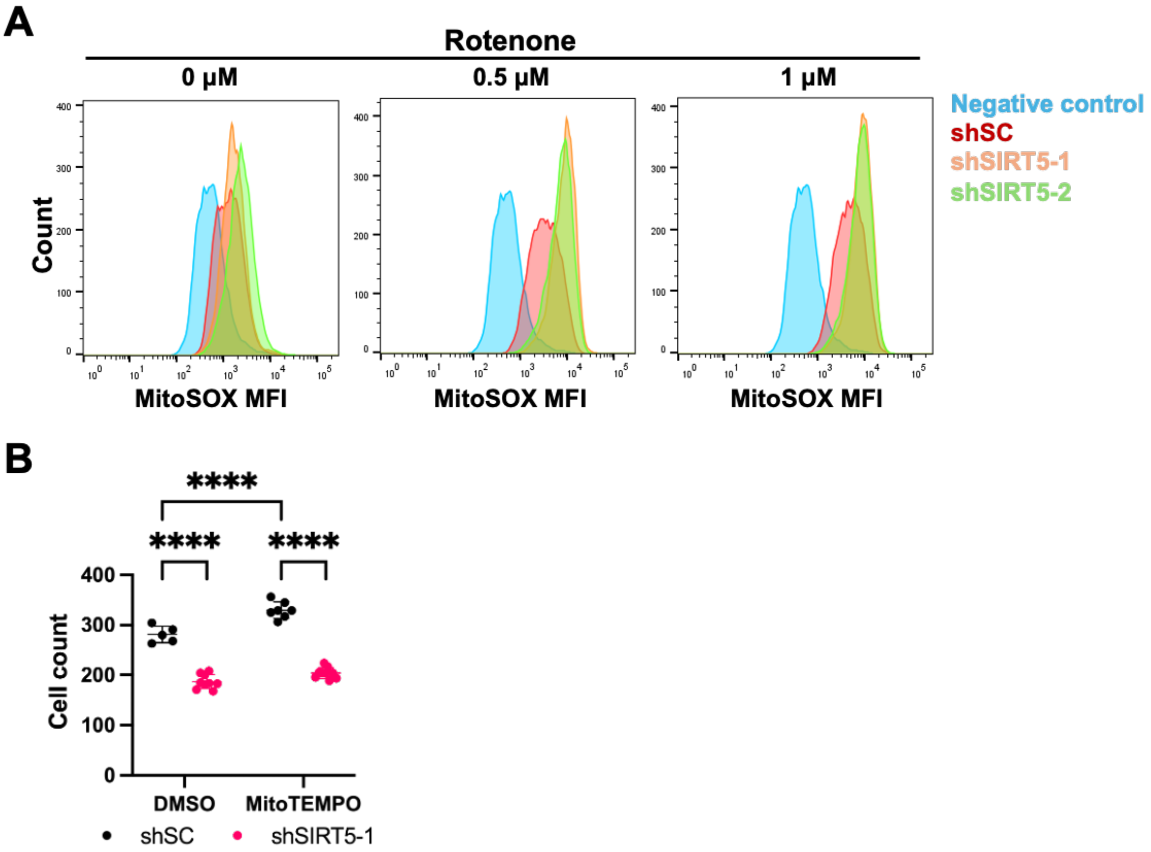


**Figure S6. *SIRT5* inhibition increases mtROS and compromises endothelial function in 3D culture. (A)** Measurements of MitoSOX MFI and cell counts in shSC and shSIRT5 HUVECs were assessed 2 hours after treatment with rotenone or DMSO. **(B)** Quantification of vessel cell number in vessels of Fig 5E (****$p < 0.001$, two-way ANOVA, n = 5-8 per group).

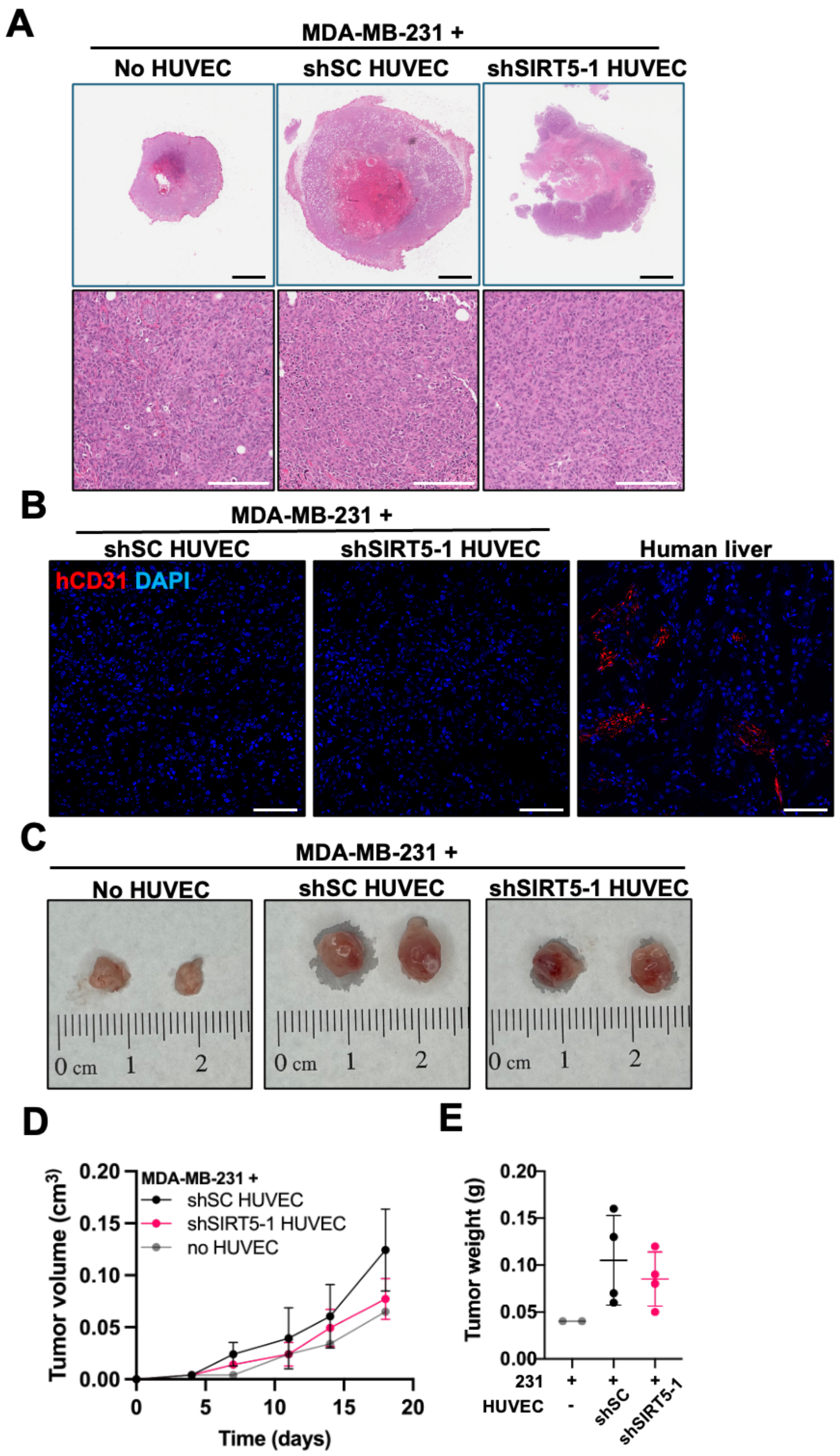
A
MDA-MB-231 +
No HUVEC
shSC HUVEC
shSIRT5-1 HUVEC
B
MDA-MB-231 +
shSC HUVEC
shSIRT5-1 HUVEC
Human liver
hCD31 DAPI
C
MDA-MB-231 +
No HUVEC
shSC HUVEC
shSIRT5-1 HUVEC
0 cm 1 2
D
MDA-MB-231 +
shSC HUVEC
shSIRT5-1 HUVEC
no HUVEC
Tumor volume (cm3)
0.00
0.05
0.10
0.15
0.20
0
5
10
15
20
Time (days)
E
Tumor weight (g)
0.00
0.05
0.10
0.15
0.20
231 + + +
HUVEC - shSC shSIRT5-1

**Figure S7. Co-injection of HUVECs enhances long-term tumor growth of transplanted 231 cells but does not persist in mice. (A)** Representative H&E images of tumor sections from NSG mice injected with MDA-MB-231 (231) cells with or without shSC or shSIRT5 HUVECs into the mammary fat pads. Scale bar (black): 2 mm. Scale bar (white): 200 µm. **(B)** Representative IF images of human CD31 (hCD31) and DAPI-stained tumors collected 35 days post-injection. Scale bar: 100 µm. **(C)** Representative images of tumors in NSG mice collected 18 days post-injection. **(D-E)** Tumor growth curves and weight at endpoints of tumors from panel (C) (two-way ANOVA, n = 2-4 per group).

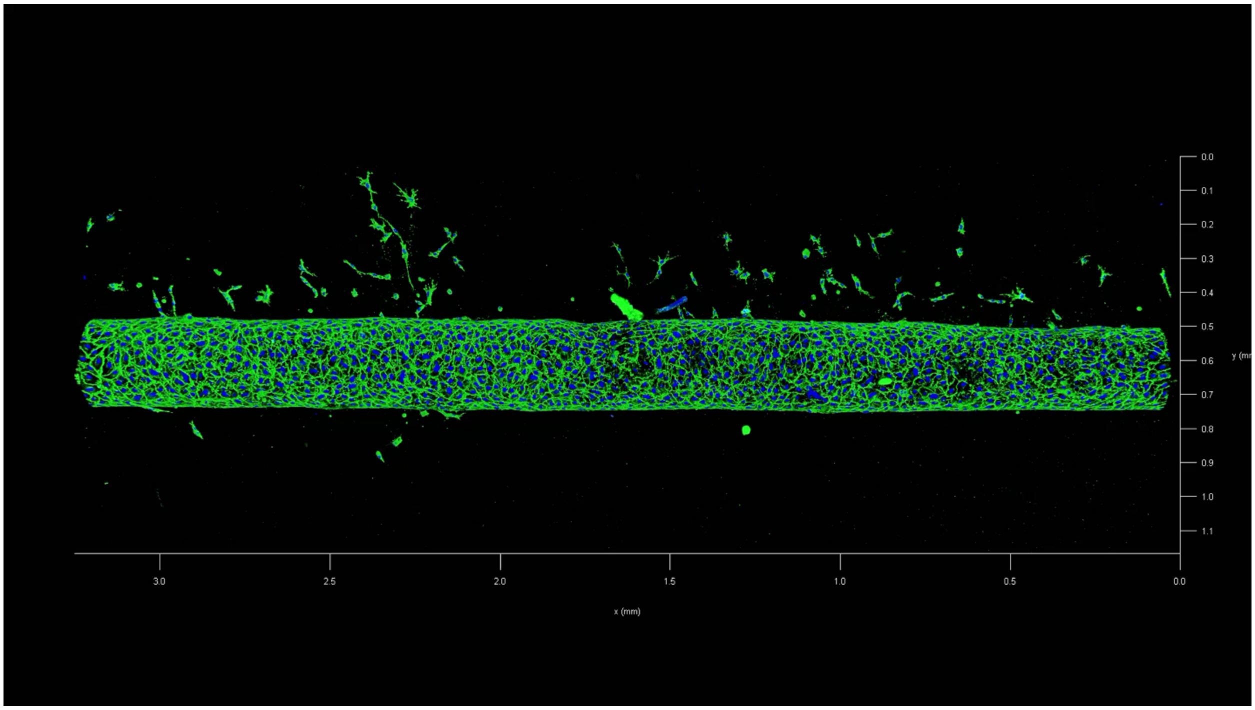


**Supplemental Video 1. 3D reconstruction of a vessel formed by HUVECs in the vessel-on-chip system.** A representative 3D rendering of a vessel-on-chip device containing HUVECs. HUVECs were fixed and IF stained for the endothelial junction marker VE-cadherin (green) and nuclei (DAPI, blue). Z-stack images were acquired using confocal microscopy and reconstructed into a 3D volume to visualize vessel architecture and lumen formation.
